# Molecular Mechanisms Driving Bistable Switch Behavior in Xylem Cell Differentiation

**DOI:** 10.1101/543983

**Authors:** Gina M Turco, Joel Rodriguez-Medina, Stefan Siebert, Diane Han, Hannah Vahldick, Christine N Shulse, Benjamin J Cole, Celina Juliano, Diane E Dickel, Michael A Savageau, Siobhan M Brady

**Author notes:** Contact Info Correspondence.

## Abstract

Plant xylem cells conduct water and mineral nutrients. Although most plant cells are totipotent, xylem cells are unusual and undergo terminal differentiation. Many genes regulating this process are well characterized, including the *VASCULAR-RELATED NAC DOMAIN7 (VND7), MYB46* and *MYB83* transcription factors which are proposed to act in interconnected feed-forward loops. Much less is known regarding the dynamic behavior underlying the terminal transition to xylem cell differentiation. Here we utilize whole root and single cell data to mathematically model this relationship. These provide evidence for *VND7* regulating bistable switching of cells in the root to a xylem cell identity, with additional features of hysteresis. We further determine that although *MYB46* responds to *VND7* induction, it is not inherently involved in executing the binary switch. A novel regulatory architecture is proposed that involves four downstream targets of VND7 that act in a cycle. These data provide an important model to study the emergent properties that may give rise to totipotency relative to terminal differentiation and reveal novel xylem cell subtypes.

## Introduction

Xylem cells are critical for the transport of water, sugars and mineral nutrients from the soil to the above ground tissues of the plant. The development of these xylem vessels and tracheary elements have facilitated the evolution of vascular plants from bryophytes, which need to grow on or near the surface of water, to trees, allowing for the transport of water over a hundred meters in distance (Raven, 1993). This essential role has led to the secondary cell wall surrounding xylem cells comprising the majority of terrestrial biomass. The xylem secondary cell wall is composed primarily of polymers; cellulose, hemicellulose and lignin, which provide the structural support needed for long-distance transport. In addition, these polymers determine both cell shape and cell specialization. Xylem vessel type can be distinguished by secondary cell wall deposition patterns. Protoxylem cells, have spiral and annular thickenings, while metataxylem cells have pitted thickenings.

In the Arabidopsis root, the xylem developmental trajectory can be separated into three distinct stages. First, xylem vessels are specified by positional signals from stem cell precursor cells (De Rybel et al., 2014). They then undergo proliferation, elongation, and finally enter the differentiation zone where they undergo secondary cell wall deposition and programmed cell death to form a long hollow vessel (Dejardin et al., 2010; Fukuda, 1996). While the majority of plant cells are totipotent, meaning that they can differentiate into any other cell type within the organism, xylem cells are unusual in that they undergo a unidirectional developmental trajectory, ending with terminal differentiation and programmed cell death. This is required for function – that is, its ability to conduct water and nutrients. This raises an important question: what molecular mechanisms facilitate the commitment to terminal cell differentiation in xylem cells, and, ultimately, how are these mechanisms distinct from the differentiation of plant cells that retain totipotency?

In contrast to plants, totipotency is not a feature of the majority of animal cells. Instead, as animal cells continue along their developmental trajectory, their ability to switch fate becomes progressively reduced. Given a quasi-steady state system **(Figure S1A**), when a cell is exposed to different concentrations of developmental stimulus, the cell can respond with two different trajectories resulting in a bistable hysteretic switch, one trajectory going from the uninduced to the induced state occurring at a high threshold and another when going from the induced to the uninduced state at a lower threshold (**Figure S1C, F**). Given the same mechanistic model (**S1A**), but without positive feedback or sufficient cooperativity, the result is a monostable-graded switch with a single trajectory when going from the uninduced to the induced state or from the induced state to the uninduced state (**Figures S1B, D, E**). Bistable hysteretic switches are common features of cell differentiation in animal cells. Examples include switching of neural progenitor cells to oligodendroglia in the brains of *Rattus norvegicus*, tracheal cell specification from a field of progenitor cells in *Drosophila melanogaster*, gut differentiation from the endoderm in *Caenorhabditis elegans*, and differentiation of the Veg_2_ lineage in *Strongylocentrotus purpuratus* embryos (Cruz-Ramirez et al., 2012; Fukushige et al., 1998; Lai et al., 2004; Maduro and Rothman, 2002; Marshall and McGhee, 2001; Metzger and Krasnow, 1999; Zelzer and Shilo, 2000; Zhu et al., 1997). We thus hypothesize that there is a transcriptional design principle (bistable-hysteresis) underlying the xylem cell developmental trajectory that initially commits these cells into this “terminal” differentiation program. A bistable hysteretic switch would facilitate this developmental commitment by creating a sharp threshold value that a developmental signal must first exceed in order to switch fates and a buffer zone that would discourage inappropriate commitment or reversal of commitment in the presence of noise.

Genes regulating xylem cell differentiation have been identified through transdifferentiation experiments. Transdifferentiation of non-xylem cells, that is any cell without xylem identity, independent of its developmental state, into xylem cells can be established through the addition of phytohormones (auxin, cytokinin and brassinosteroids) (Fukuda, 1992; Fukuda and Komamine, 1980; Roberts, 1976; Sachs, 1968). Alternatively, xylem transdifferentiation can be induced by the glycogen shaggy kinase IIIs inhibitor, bikinin (Kondo et al., 2015). A hormone-mediated transdifferentiation study led to the identification of the transcription factors *VASCULAR-RELATED NAC-DOMAIN PROTEINS 6 and 7 (VND6 and VND7)* as being sufficient to drive xylem cell differentiation (Kondo et al., 2015; Kubo et al., 2005; Yamaguchi et al., 2010) and thus these genes are considered as master regulators of xylem differentiation. Transdifferentiation studies were also used to identify direct downstream targets of VND7; the transcription factors *MYB83* and *MYB46* are suggested to act redundantly in the second tier of a transcriptional cascade controlling xylem secondary cell wall biosynthesis (McCarthy et al., 2009; Yamaguchi et al., 2011) through a series of feed-forward loops (FFL) (**Figure 1**) (Mangan and Alon, 2003; Mangan et al., 2003; McCarthy et al., 2009; Taylor-Teeples et al., 2015). In plants, the potential for an asymmetric cell division state was linked via mathematical modeling to bistability of nuclear localized SHORT-ROOT and SCARECROW transcription factor levels generated through feedback within FFLs (Cruz-Ramirez et al., 2012). We thus hypothesize that VND7 stimulates or participates in the commitment to xylem cell differentiation via a bistable hysteretic switch through these FFLs.

**Figure 1.**
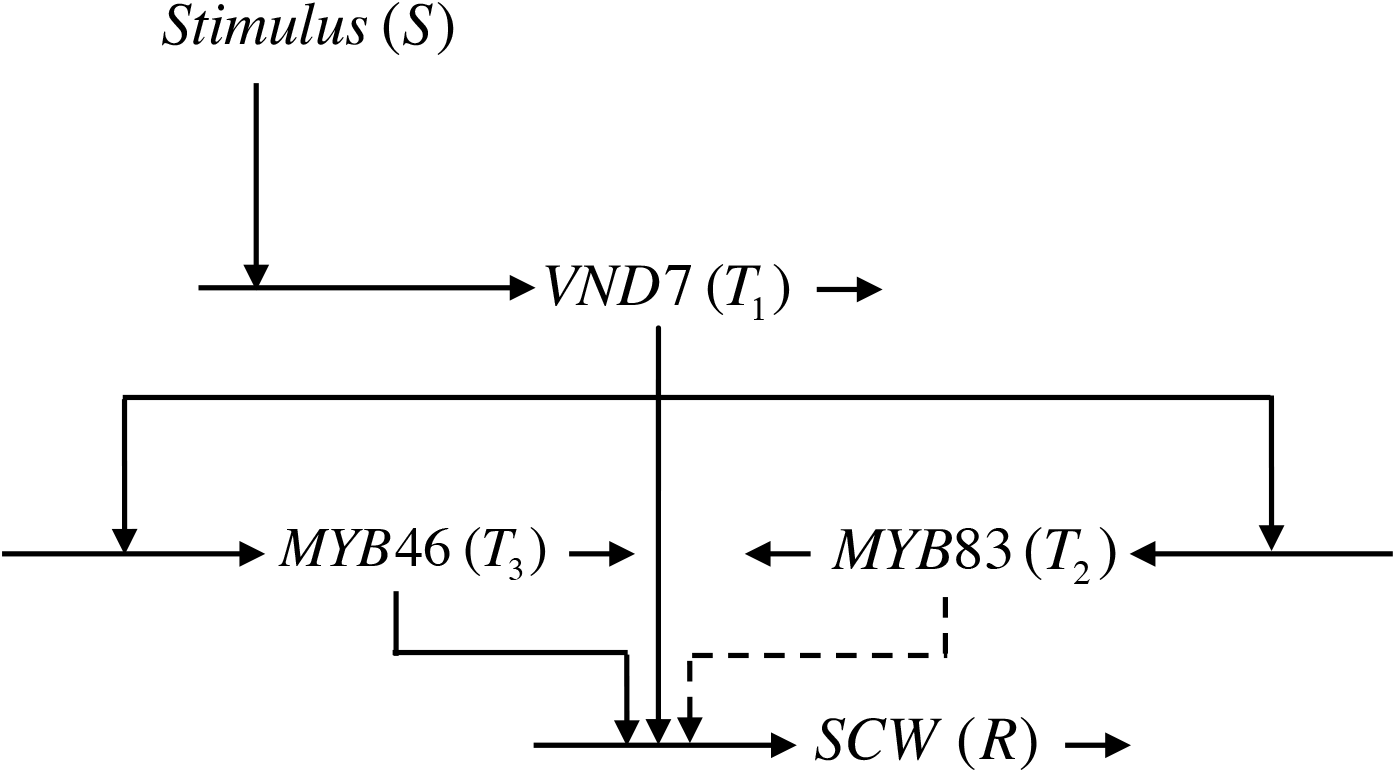
Predicted feed-forward loop topology for xylem differentiation. Here VND7 is turned on in response to a stimulus (S) where solid lines indicate known binding and regulation of a transcription factor (T_i_ ) to the promoter region of indicated targets (R) (marked by the directionality of arrow) while dotted lines represent inferred interactions.

Here, we first provide evidence for a bistable switch with features of hysteresis in whole roots and in single cell transcriptome data. We then use these single cell sequencing data and mathematical modeling to determine that neither MYB46 nor MYB83 respond to VND7 in a switch like fashion, and identify gene targets that respond in a monostable or bistable switch-like fashion during xylem differentiation. Further, a putative circuit is predicted for the bistable switch and potential interactions and parameter configurations where this feedback can generate bistability are suggested. Together, these data provide insight into a design principle that underlies the formation of a cell type critical for plant growth, development and the transition of plants from water to land.

## Results

### The relationship between *VND7* expression and xylem formation resembles a bistable switch

To determine if VND7 regulates xylem differentiation via a bistable or monstable switch, we used β-estradiol-*VND7* inducible transgenic Arabidopsis plants (Coego et al., 2014). This two-component system drives expression of *VND7* in all cell types under the G10-90 constitutive promoter (Zuo et al., 2000). We varied the expression of *VND7* in a dose-dependent manner (using estradiol as the inducer) and then quantified the number of xylem cells in the elongation zone of the root where protoxylem differentiation typically occurs (**Figure 2A-B**) (Benfey and Scheres, 2000; Esau, 1977). Roots were stained with Calcofluor white to stain for cellulose and with basic fuchsin to mark lignin deposition, both a hallmark of xylem differentiation associated with secondary cell wall biosynthesis (**Figure 2B)** (Ursache et al., 2018). The ratio of cells with spiral patterning to the total number of cells in each root image was used as a measure of percentage transdifferentiation to xylem and ranged from 0-100% (**Figure 2B-C, Table S1**). In plotting the resulting data, we make the assumption that each cell within the root (the population of cells) encounters approximately the same concentration of β-estradiol and thus, *VND7* expression. When *VND7* expression was increased approximately 5-fold, a dramatic jump in xylem cell differentiation was observed. Cell types closer to the root vascular tissue were much more likely to transdifferentiate into xylem cell types upon β-estradiol treatment, suggesting these cells may have more plasticity in their response to *VND7* induction (**Figure S2**).

**Figure 2.**
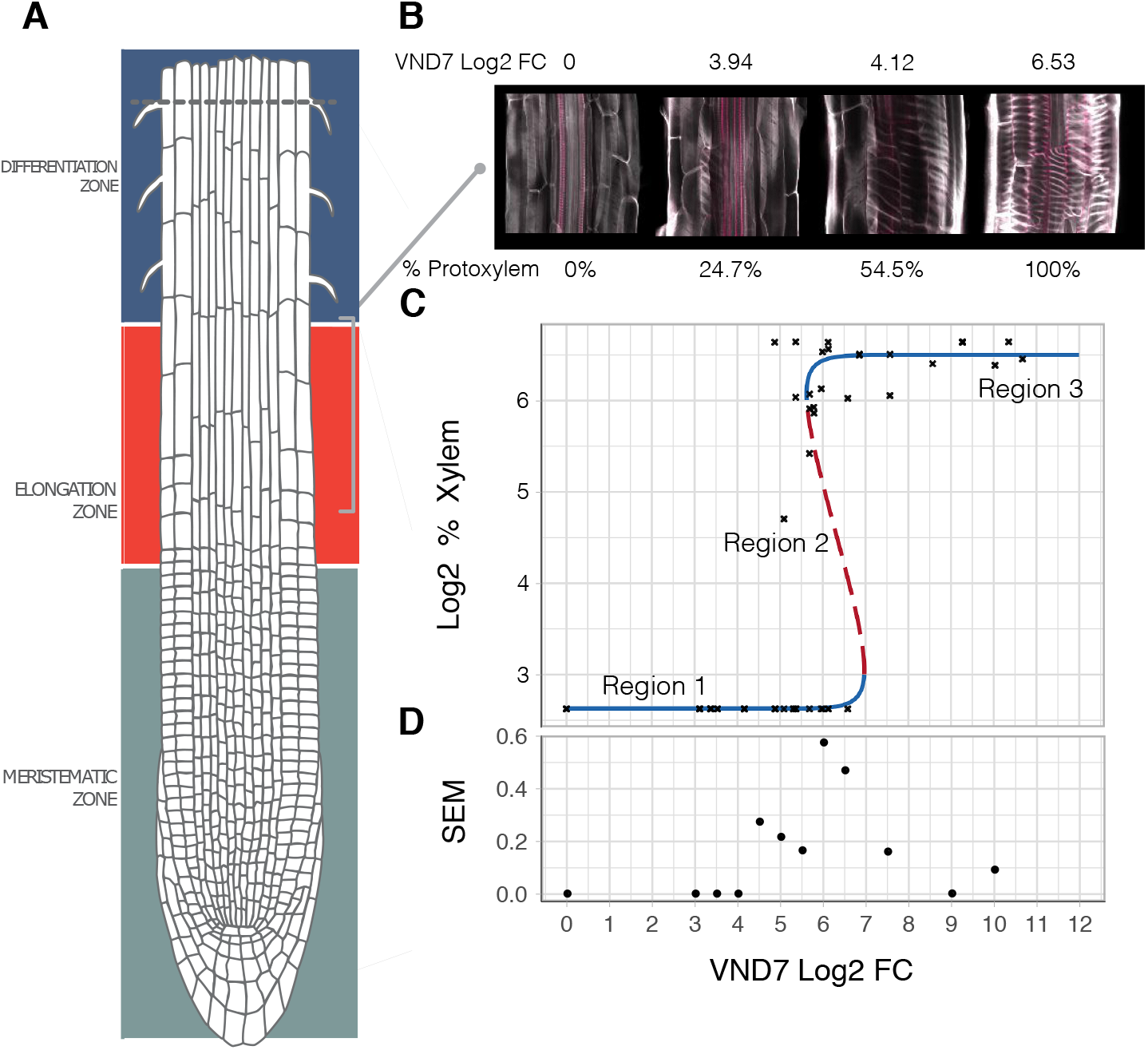
*VND7* expression regulates xylem formation as a bistable switch with features of hysteresis. **(A)** Xylem cell differentiation was measured in the root elongation zone. Image of Arabidopsis root:(Frédéric, 2017). **(B)** Differentiation was measured using calcofluor-white and fuchsin to image cellulose (white) and lignin (pink) as hallmarks of xylem cell differentiation. **(C)** Each X-mark represents an individual root’s xylem phenotype and its estimated VND7 FC, inferred from RNA extracted in bulk from the plate of origin. Plates are grown at varying estradiol concentrations each containing at least three biological replicates (separate plates). The % of xylem cells is represented in log format for a log-log comparison to *VND7* log fold change in response β-estradiol induction, the best fit to the data is provided by the generic model in Methods with the following values for the parameters: *P*_*max*_= 90.6 (% xylem cells), *P*_*min*_= 6.1 (% xylem cells under normal conditions), *K*_*S*_= 10 (*VND7* FC), *K*_*p*_ = 260 (*VND7* FC), m =4, and n = 4. Stable state = solid line, unstable state = dashed line. **(D)** To plot variance as a function of *VND7* FC, the <u>s</u>tandard <u>e</u>rror of the <u>m</u>ean (SEM) was calculated for each *VND7* log fold change (n data-points >10).

The observed relationship between *VND7* and xylem differentiation (**Figure 2C**) suggests the existence of a bistable switch with three distinct regions: (1) a region of monostable minimal (wild type) levels of xylem cell differentiation when the value of *VND7* expression is below a lower threshold (5-fold change, Region 1), (2) a region of fully induced monostable xylem differentiation, when the value of *VND7* expression is above an upper threshold (7-fold change, Region 3), and (3) a bistable region in which xylem differentiation is either at its maximum or minimum, when *VND7* expression is between the upper and lower switch thresholds of *VND7* expression (between 5-and 7-fold change, Region 2). The two stable states are separated by an unstable state from which xylem differentiation diverges to the minimal or maximal levels of xylem differentiation. A *Separation Score* was used to determine the existence of two stable states (see Methods). A separation score nearer to 0.3 is indicative of a uniform distribution of data and thus only a single/mono stable state. At low expression of *VND7* (1 to 4 FC), the separation score was 0, at moderate *VND7* expression (4 to 7 FC) the separation score was 0.82, while at high *VND7* expression (7-10 FC), the separation score was 0.4 (**Figure 2C**).

Using the mechanistic model described in **Figure S1A**, we next fit the parameters obtained in **Figure 2C**, to determine the values of the parameters consistent with the bistable commitment to differentiation in response to the induction of VND7. Given that this model is for a single cell, we carried out stochastic simulations to determine how a monostable (**Figure S3A-G**) or a bistable (**Figure S3H-N**) switch would behave under different levels of stimulus (equivalent to VND7 induction). In these simulations, ON is equivalent to a commitment to terminal differentiation, and OFF is the absence of this commitment. The simulated responses in **Figure S3** (red and green lines) are representative of numerous simulations that give very similar results. The monostable system has only a single mean behavior at all levels of input simulation (**Figure S3A-G**). In contrast, the bistable hysteretic response (**Figure S3H-N**) only shows a single mean behavior below a lower threshold (**Figure S3N**) and above an upper threshold (**Figure S3K, L**) of input stimulation; whereas there are two distinct mean behaviors with one ON and the other OFF at intermediate levels of input stimulation (**Figure S3J**). In the ON state (**Figure S3L**), the bistable system only returns to the OFF state when the stimulus is reduced to a level less than the lower of the two thresholds (**Figure S3N)**. Thus, in the context of the cell population, only the bistable system is consistent with the data presented in **Figure 2C**.

An additional characteristic of bistability with hysteresis is increased variance marking the unstable region (Region 2). We used a sliding window to determine the variance in xylem formation via standard error of the mean (SEM) for each binned *VND7* fold change expression value (**Figure 2D**). A sharp increase in variance was observed within the predicted thresholds for switching between 5 and 7-fold change (**Figure 2D**). This suggests discontinuous behavior, which also tends to support the presence of bistable-hysteresis. Together, these data provide evidence that VND7 is a sufficient input stimulus for xylem differentiation as a bistable switch with features of hysteresis.

### Single cell sequencing reveals a dramatic pattern of bistable switching to xylem cell identity in response to *VND7* expression

Transdifferentiation is the process by which differentiated cells transform into a completely different cell type. In the root elongation zone, we observed evidence of a bistable-hysteretic switch amongst the population of cells within the elongation zone. To test how *VND7* changes the transcriptional and developmental trajectory of individual cells with different identities, we carried out single cell RNA-seq profiling using Drop-seq at the minimum/uninduced (0 µM β-estradiol) and at the maximum/induced (20 µM β-estradiol) concentrations (Macosko et al., 2015), where 100% conversion was observed (**Figure 2C**). Two hundred and fifty cells were obtained, before applying a minimum cutoff for the number of detected genes, from DropSeq for each of the two conditions (**Figure 3A, Table S2)** (Kondo et al., 2015; Yamaguchi et al., 2010) and three hundred and seventy-four transcriptomes obtained. Cell identity was determined in an unsupervised manner and then compared between the minimum and maximum *VND7* expression levels to determine how VND7 influences the transformation of cell identity. To determine identity, cells were clustered using a shared nearest neighbor-based algorithm in Seurat (Satija et al., 2015). The identity of the eleven resulting clusters (**Figure 3A**) was determined by their average gene expression profiles using a predefined Index of Cell Identity (ICI) algorithm from Arabidopsis roots (**Figure 3A-B**) (Birnbaum et al., 2003; Birnbaum and Kussell, 2011; Brady et al., 2007; Dinneny et al., 2008; Efroni et al., 2015; Gifford et al., 2008; Lee et al., 2006; Nawy et al., 2005). As expected, a significant increase in the number of maturing xylem cells (protoxylem identity) was observed in the *VND7*-induced cell population (adjusted p<2.2×10^−16^). A less dramatic, although significant increase was also observed in the number of phloem cells as previously seen (adjusted p=0.0002) (**Figure 3A-B)** (Kondo et al., 2015; Kondo et al., 2016). In comparison, the numbers of cortex, trichoblast and endodermis cells were significantly reduced in the 20 µM β-estradiol-treated cells relative to the uninduced cells (**Figure 3A-B**). Based on this change in cell proportions, we can infer that cortex, trichoblast and endodermis cells are more likely to transform into differentiating xylem cells. This would need to be confirmed by measurement of *VND7* expression or protein levels in these cell types after induction.

**Figure 3.**
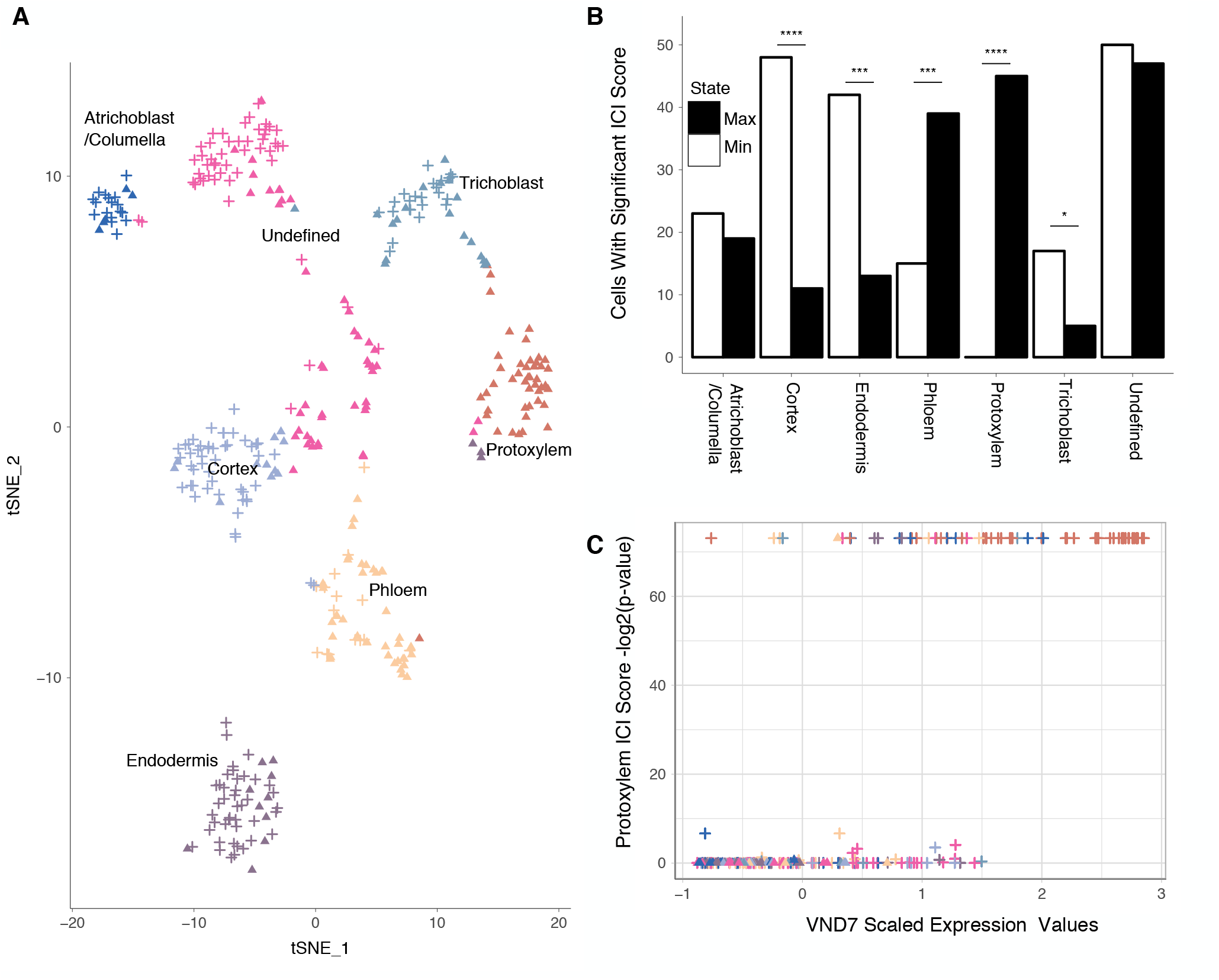
Evidence for a bistable transcriptional switch in single cell sequencing profiles. **(A)** t-SNE clustering of 374 single cells from Arabidopsis roots either over-expressing *VND7* (triangle) or with wild type levels of *VND7* expression (+). Clusters are labeled according to their cell identity score (ICI). **(B)** Quantification of cell type with cell identity determined from **A** in the induced population (20 *µ*M β-estradiol; denoted as VND7 over-expression OE), compared to the total number of cells with that identity in the uninduced cells, (0 *µ*M β-estradiol; denoted as control). **** represents an adjusted p-value ≤ 0.0001, *** p-value ≤ 0.001, * p-value ≤ 0.05 as determined by a Fisher’s-Exact test with Benjamini-Hochberg correction. **(C)** Significant protoxylem ICI scores determined for individual cells for induced (+) and uninduced (triangle) populations, plotted against *VND7* normalized expression data.

The single cell transcriptomes of these 374 cells provide a higher resolution (relative to whole root expression profiling) to quantify the breakpoints between the two stable states in relation to *VND7* expression. Using this increased power to quantify the cell identity switch, we calculated the ICI score for each individual cell’s transcriptome (**Figure 3C, S4**). We then plotted *VND7* gene expression relative to the p-value associated with an individual xylem cell ICI score (**Figure 3C**). The number of cells with a significant protoxylem ICI score is very low for normalized *VND7* levels below a relative log2 value of 2, either very low or very high for *VND7* levels between 2 and 3.5, and very high for *VND7* levels above 3.5, consistent with a bistable switch (**Figure 3C**). A similar switch was observed in the “native” context using 12,000 Col-0 single cell sequences, (Shulse et al., 2018) (**Figure S5A**) and in “native” xylem temporal data of the Arabidopsis root (Cartwright et al., 2009) (**Figure S5B**). The separation scores support a clear bistable pattern in the single cell data: the scores for low (0-2) and high (4-6) *VND7* expression levels was 0, while for mid *VND7* levels (4-6), the score was 0.98. Thus, within every measured cell within the Arabidopsis root, *VND7*-mediated xylem cell transdifferentiation is consistent with a bistable switch as observed in the transcriptional profiles to a xylem identity.

### Known VND7 targets and their association with the bistable switch

If VND7 stimulates xylem differentiation as a bistable switch, then we expect to also observe bistable regulation of downstream targets as an emergent property of the network. Depending on the network topology we may also observe a subset of VND7 targets with a monostable response, as these are not mutually exclusive (described in **Figure S1D)**. To determine the downstream targets exhibiting a switch-like behavior we took advantage of the single cell variance in *VND7* expression. The distribution of *VND7* expression levels across cells is not bimodal (**Figure S4D**) and its separation score in induced cells with a xylem identity is 0.34 which is the expected for a uniformly random response (our null hypothesis). Thus, there is no evidence of a sharp increase in *VND7* nor its participation in a positive feedback loop generating bistability. We used these data to examine the relationship between *VND7* and potential target genes among 203 correlated genes (r^2^>0.5) to identify monostable or bistable expression patterns.

Given these three expression bins, (low, mid-and high), 116 of the 203 positively correlated genes had significantly different VND7-induction at mid-and high-levels of induction relative to that in un-induced cells. If a gene was responding to VND7 induction in a monostable fashion, target gene expression should increase in a graded manner concomitant with an increase in VND7 expression. In a bistable case, the target gene should change its expression suddenly and in a dramatic fashion between the low and mid-levels of VND7 expression, but there should be little difference in the expression of the target gene between the mid-and high VND7 expression values (Methods). 161 genes changed their expression in a monostable fashion as determined by a statistically significant difference between the middle and high expression bins (**Table S4**). In comparison, 15 genes showed a dramatic increase in expression between the low and middle VND7 expression bins, but no statistically significant change between the middle and high bins, and are thus candidates for genes that participate directly in the bistable switch (**Figure S6, Table S4**). No defined GO terms were significantly over-represented in the bistable switch responsive category. GO terms associated with SCW synthesis were identified for the monostable switch categories (**Table S4**). Neither VND7-, MYB46-nor VND7/MYB46 joint direct targets were enriched in the bistable gene set (Kim et al., 2013; Ko et al., 2009; Yamaguchi et al., 2010; Yamaguchi et al., 2011). Given that MYB46 expression is not bistable, and the lack of enrichment of their direct targets in the bistable gene set, the VND7-MYB46 FFL is not directly involved in generating the bistable switch observed. Furthermore, given that MYB83 expression is not induced in response to VND7 induction (greater than r=.5), the VND7-MYB83 FFL is also likely not involved in xylem cell transdifferentiation.

### Single cell sequencing identifies cohorts of cells that are targets of VND7, MYB46 and MYB83-dependent vascular processes

MYB83 is sufficient to drive lignin biosynthesis associated with the xylem secondary cell wall and shares many targets with *MYB46* (McCarthy et al., 2009). These data suggest that MYB83 acts in a partially redundant manner to MYB46 and in a parallel feedforward loop. However, unlike *MYB46, MYB83* expression is not induced upon *VND7* induction (**Tables S4**). To further explore the role of these transcription factors in VND7-dependent xylem transdifferentiation, we carried out Drop-seq experiments on *MYB46* and *MYB83* β-estradiol inducible plants. The transcriptomes of 776 *MYB46*-induced cells and 989 *MYB83*-induced cells were collected (**Table S2**), and the extent of transcriptome variation was determined by combining the *MYB46, MYB83* and *VND7*-induced cell transcriptomes and their corresponding controls in Seurat (**Figure 4A**). Within the eleven resulting clusters, three clusters with a high protoxylem ICI score were identified (**Figure 4A-B, Figure S7**). To assess if these clusters could be used to differentiate VND7-MYB46-FFL and VND7-MYB83-FFL regulation, we quantified the number of cells originating from each overexpression experiment within the three protoxylem clusters (**Figure 4C**). Clusters were labeled accordingly; mixed (Cluster 7, contains induced and uninduced cells from *VND7, MYB83* and *MYB46* over-expression experiments), *VND7* primary (Cluster 10) or *MYB46* primary (Cluster 5) (**Figure 4C**). Single cell sequencing has been used to classify novel cell subtypes (Macosko et al., 2015), and we propose that these three clusters represent subtypes of xylem cells. To confirm that these xylem subtypes occur in natural cell populations we included 16 “native” xylem cell transcriptomes from Arabidopsis root cells (Shulse et al., BioRXIV, 2018). Each of our three-xylem clusters contained cells from “native” cell populations (**Figure S7C-D**). To determine the molecular function of each cluster/cell subtype, a range of xylem-related biological process gene sets were determined for statistically significant over-representation (**Figure 4D**, **Table S5**).

**Figure 4.**
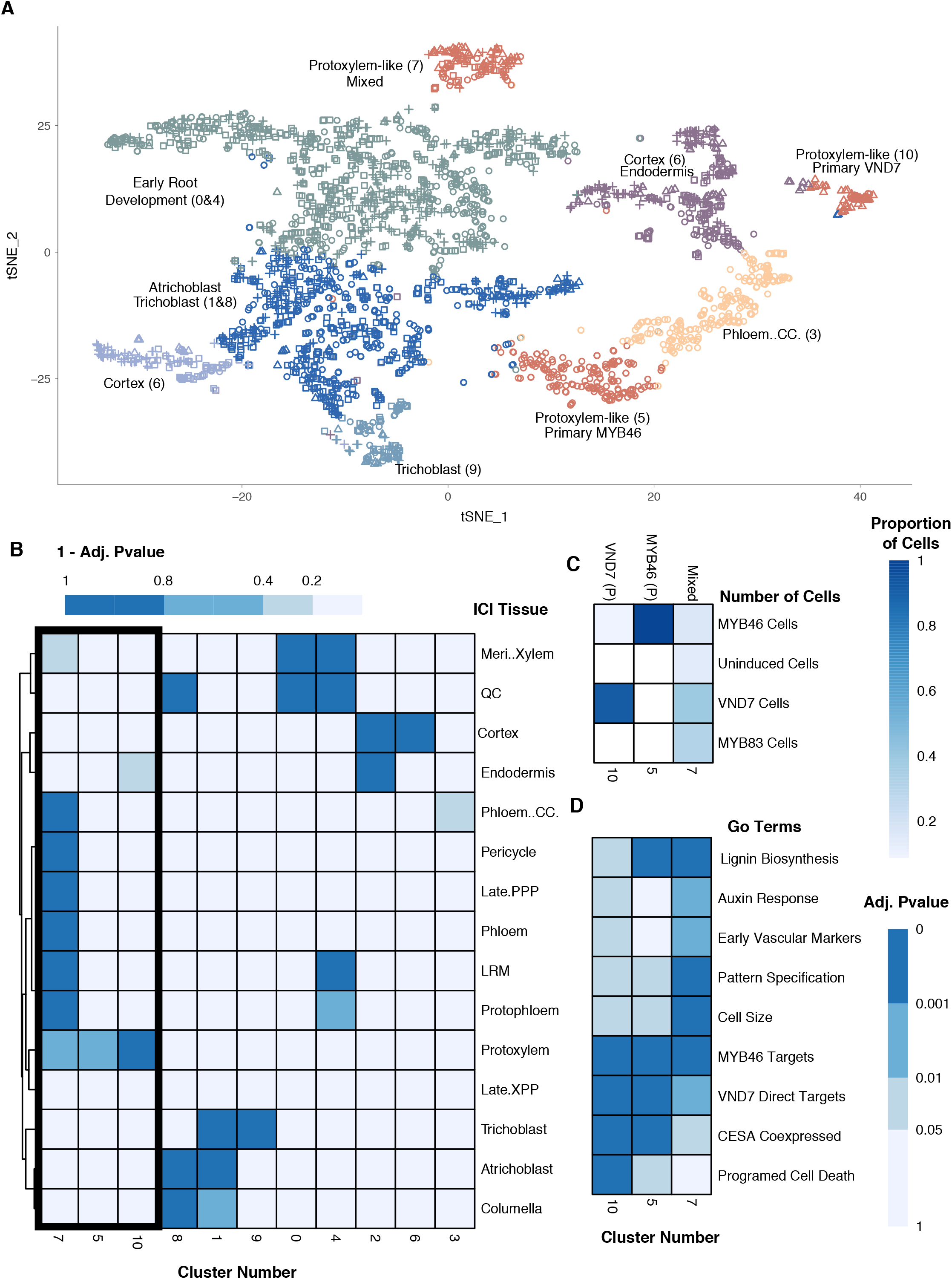
Single cell transcriptomes overexpressing *VND7, MYB46* and *MYB83* separate distinct vascular developmental processes. **(A)** t-SNE clustering of single cells from all Arabidopsis root overexpression experiments for either *VND7* (triangle), MYB46 (circle), MYB83 (square) or uninduced cells (+). Clusters are labeled according to their cell identity score (ICI) and corresponding cluster number. **(B)** Heatmap showing enrichment for ICI scores within each cluster, with the three protoxylem clusters highlighted. **(C)** The three protoxylem clusters and the proportion of cells within each cluster that correspond to each overexpression experiment, cluster numbers are the same as those in **(D)**. White boxes indicate no cells are present for the corresponding experiment. Clusters were also labeled based on their experimental distribution as mixed, VND7 P. (*VND7* Primary) or MYB46 P. (*MYB46* Primary). **(D)** Enrichment of xylem-related genes within the three protoxylem clusters.

The *VND7*-primary subtype contains an over-representation of genes associated with cells undergoing late stages of xylem development, including programmed cell death marker genes, bistable genes, protoxylem cell identity genes (**Figure 4B-C**) and genes defined by the GO terms “cell wall polysaccharides” and “xylem” as well as enrichment for genes displaying “monostable” behavior (**Table S4 and S5**). The *MYB46*-primary cluster showed similar patterns of enrichment for programmed cell death marker genes and bistable genes, but also a more extreme enrichment in genes associated with lignin biosynthesis than the VND7 primary cluster (**Figure 4D**). The *MYB46*-primary cluster differs from the “mixed” and the *VND7*-primary cluster in the absence of enrichment of developmental genes, including auxin-responsive genes and genes associated with cell expansion (**Figure 4D**). In contrast, genes of the “mixed” cluster, showed significant enrichment in developmental categories such as auxin-responsive (including *PIN3* and *PIN7*), cell expansion and pattern specification (**Figure 4D**), and significant depletion for the “monostable” gene set (**Table S4**). In addition, a significant increase in expression was observed for the early vascular development genes *TMO5* and *WOL* in the “mixed” cluster (Cano-Delgado et al., 2010; Cano-Delgado et al., 2000; De Rybel et al., 2013; Mahonen et al., 2000). Together these distinct molecular signatures reflect unique aspects of transcription factor-mediated regulation of vascular cell development and secondary cell wall biosynthesis.

### Network topology for xylem cell differentiation

The morphological and molecular data presented here demonstrates that xylem cell differentiation occurs via a bistable switch-like mechanism stimulated by VND7 but not MYB46 or MYB83. The single cell transcriptome data provides an opportunity to identify the network topology needed for xylem cell differentiation. Given that *VND7* and *MYB46* expression both display monostable expression patterns, they provide an input stimulus to but do not participate in the positive feedback loop responsible for the bistable expression of their target genes. Of these 15 bistable genes, 4 are VND7 direct targets (Class A), 1 is a MYB46 direct target (Class C) and 10 are neither VND7 or MYB46 direct targets (Class B) (**Table S4, Figure 5A).** Thus, expression of a set of hypothetical genes “X” must respond to VND7 induction and provide the intermediary link to the 15 bistable genes.

**Figure 5.**
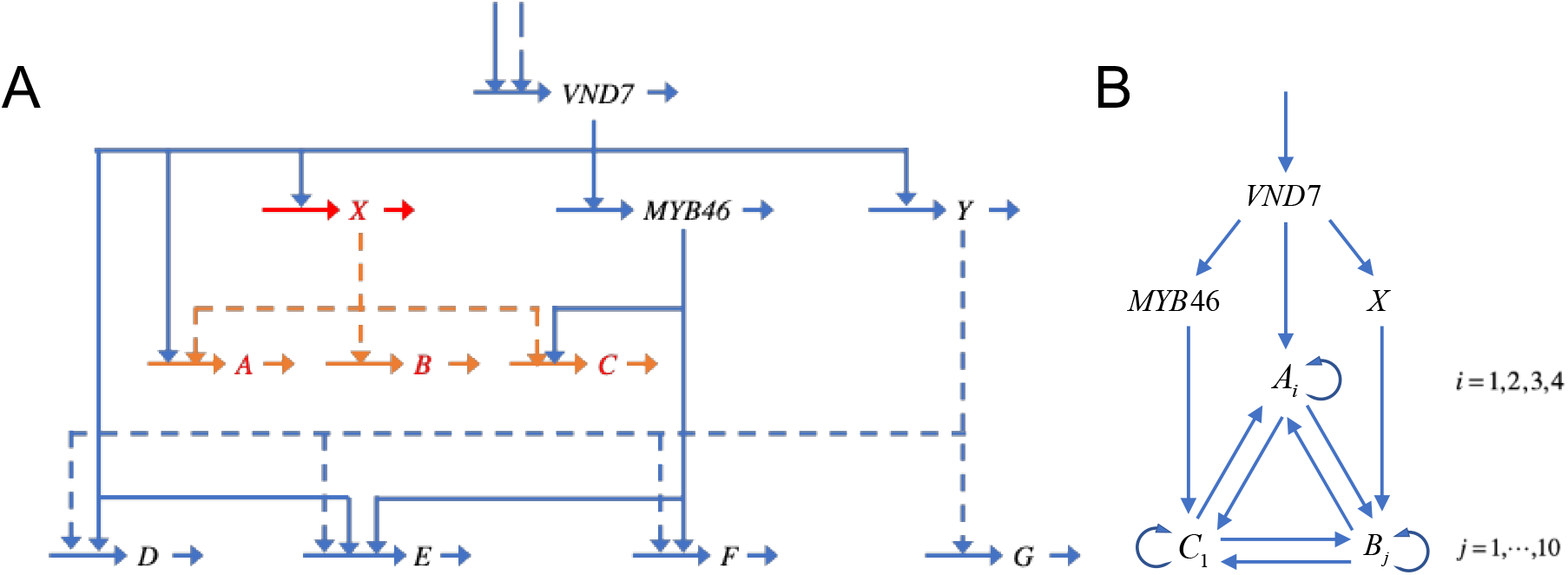
Classes of gene circuitry identified from single cell transcription data. (A) Classes of downstream target genes: **A-G**. Classes of bistable genes: A (4 VND7 direct targets), B (10 VND7 indirect targets), and C (1 MYB46 direct target). Classes of monostable genes: D (13 VND7 direct targets), E (5 VND7 & MYB46 direct targets), F (24 MYB46 direct targets), and G (37 VND7 indirect targets). Potential intermediate genes: X (targets of VND7, signals for downstream bistable genes) and Y (targets of VND7, signals for downstream monostable genes**). (B) Genes exhibiting bistability with possible interactions capable of generating the necessary positive feedback.** There are four genes (*A*_*i*_) in the class of VND7 direct targets, ten genes (*B*_*j*_) in the class of VND7 indirect targets, and one gene (*C*_*1*_) in the class of MYB46 direct targets.

Class C contains one gene with a MYB46 binding site, but no VND7 binding site, in its regulatory region (**Figure 5B**). Given that *MYB46* itself responds in a monostable fashion, the bistable response of the single gene in this class must involve autoactivating of its own promoter or receiving input from another bistable gene in Class A or B **(Figure 5B).** Class B contains 10 genes, which generates too many possibilities to examine with the resolution of data available. Thus, we carried out a more detailed analysis of the four Class A genes to predict which regulatory architectures could most likely result in the bistable switch observed.

To refine the potential circuitry of the 4 bistable genes directly regulated by VND7 (Class A) we used the phenotype-centric modeling strategy and system design space analysis to predict circuity, parameter values and robustness of phenotypes (Fasani and Savageau, 2010; Lomnitz and Savageau, 2013, 2015, Valderrama-Gómez, et al, 2018) suggested by the single cell induction data. Given that VND7 is the stimulus for the bistable switch, we focused on generic circuitry involving the four target genes that are also targets of VND7, as shown in **Figure 6A**. The phenotype-centric approach represents this mechanism in terms of kinetic equations (Methods) that are used to predict qualitative patterns of dynamic expression, referred to as “phenotypes” (Savageau et al., 2009). An expression “phenotype” could include oscillatory, monostable graded or bistable hysteretic switch responses. A necessary condition for bistability is the overlap of three elemental phenotypes (**Figure S1F**) – two stable phenotypes separated by one that is unstable. The unstable phenotype must have one eigenvalue with a positive real part, indicative of an exponential instability that drives the system toward fully OFF or fully ON expression, and the expression must *decrease* in response to an *increase* in the stimulus (i.e., the logarithmic gain must be negative).

**Figure 6.**
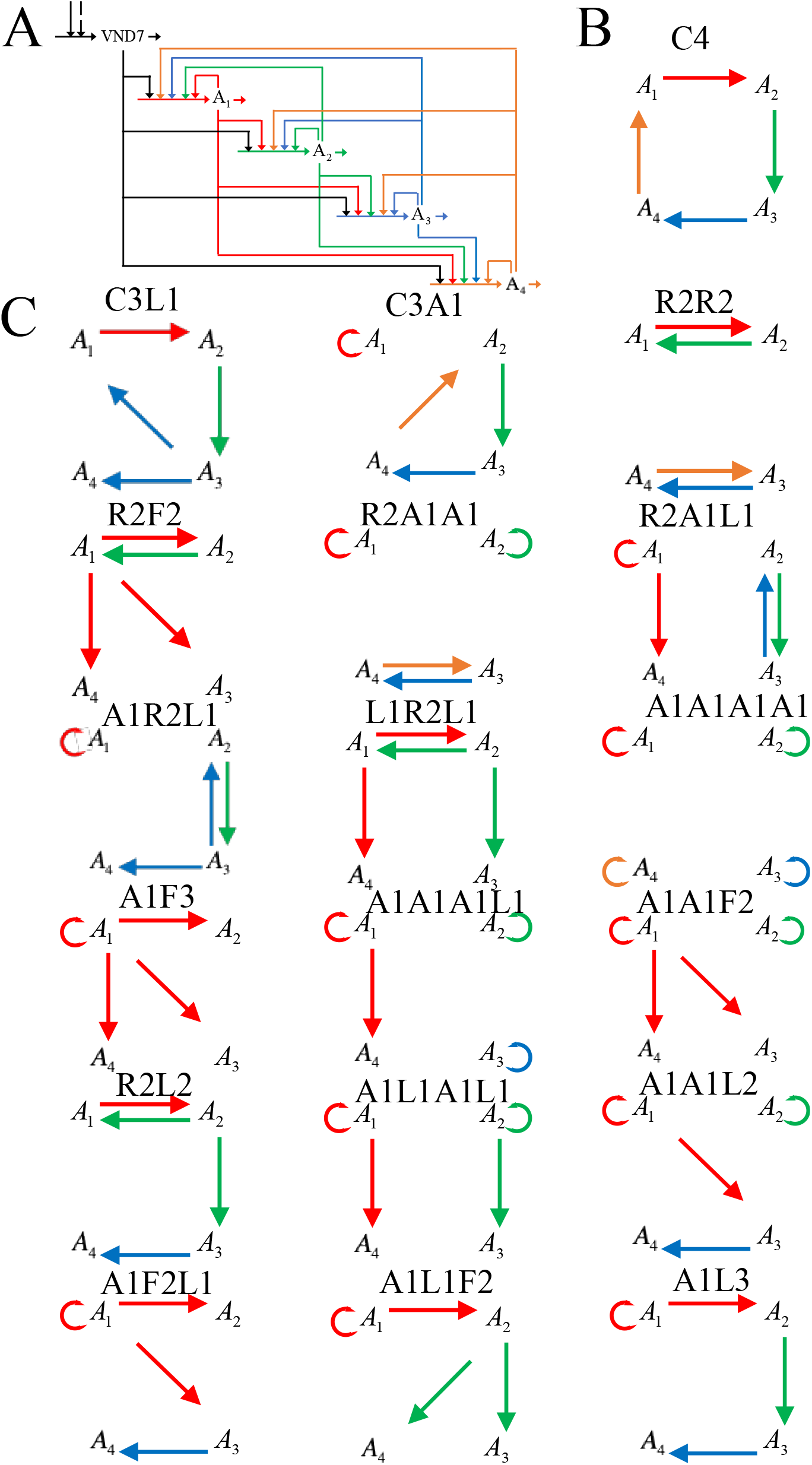
Generic circuitry for Class A bistable genes. **(A)** Circuitry showing activators binding to promoter targets and degradation of regulators. **(B)** Cartoon emphasizing only the topology of the regulatory circuit with the greatest Global Tolerances. **(C)** Cartoon circuitry for the other 18 phenotypes with the positive feedback capable of generating bistability. The circuits are ranked from left to right and top to bottom according to their Global Tolerances (**Table 1**).

The first step in the phenotype-centric modeling strategy is to enumerate the phenotypic repertoire of the model. The generic circuitry in **Figure 6A** has a total of 6,561 phenotypes. We next filtered this list to obtain bistable phenotypes that have: (a) an eigenvalue with a positive real part, indicative of an exponential instability that drives the system toward fully OFF or fully ON expression of all four genes, (b) expression of all four genes decrease in response to an increase in VND7 within the unstable region (all four logarithmic gains must negative), and (c) maximum levels of expression that differ by less than 10 fold among the four genes. A smaller subset of 256 phenotypes meet these three criteria. Given the 4-fold rotational symmetry of this circuit, many of these phenotypes are identical. The properties of the different phenotypes are listed in **Table 1**.

**Table 1.** Phenotypes of the circuitry in Figure 6A that are capable of generating a bistable induction. Phenotypes were determined using the System Design Space Toolbox. There is some combination of values for the parameters of the model that guarantees the existence of each predicted phenotype.

| Topology of Activations<sup>†</sup> | Number of Phenotypes* | Representative Phenotype* | Volume in Design Space | Global Tolerance<sup>§</sup> |  |  |
| --- | --- | --- | --- | --- | --- | --- |
|  |  |  |  | Minimum | Maximum | Mean |
| C4 | 6 | 336251 | 4.79E+18 | 10.0 | 75.0 | 27.4 |
| C3L1 | 24 | 258126 | 2.52E+15 | 4.69 | 56.2 | 15.3 |
| C3A1 | 8 | 148751 | 1.15E+15 | 3.16 | 56.2 | 14.4 |
| R2R2 | 3 | 186251 | 3.17E+14 | 5.62 | 31.6 | 13.0 |
| R2F2 | 12 | 175626 | 5.20E+13 | 3.88 | 31.7 | 11.4 |
| R2A1A1 | 6 | 123751 | 1.79E+12 | 3.16 | 31.6 | 8.76 |
| R2A1L1 | 12 | 131876 | 1.78E+12 | 2.37 | 31.6 | 8.76 |
| L1R2L1 | 12 | 178751 | 8.55E+11 | 3.88 | 31.7 | 8.28 |
| A1F3 | 4 | 97501 | 8.25E+11 | 2.61 | 31.6 | 8.25 |
| A1R2L1 | 24 | 133126 | 7.53E+11 | 3.16 | 31.6 | 8.20 |
| A1A1F2 | 12 | 113126 | 5.62E+10 | 2.37 | 31.6 | 6.71 |
| A1A1A1A1 | 1 | 121251 | 1.00E+10 | 3.16 | 10.0 | 5.88 |
| A1A1A1L1 | 12 | 119376 | 1.00E+10 | 2.37 | 31.6 | 5.88 |
| A1L1A1L1 | 12 | 116251 | 1.00E+10 | 2.37 | 31.6 | 5.88 |
| A1F2L1 | 24 | 98751 | 2.10E+09 | 1.57 | 31.6 | 5.21 |
| A1L1F2 | 12 | 101251 | 2.10E+09 | 1.57 | 31.7 | 5.21 |
| R2L2 | 24 | 180001 | 1.37E+09 | 1.94 | 31.6 | 5.04 |
| A1A1L2 | 24 | 114376 | 3.16E+08 | 1.54 | 31.6 | 4.51 |
| A1L3 | 24 | 101876 | 4.37E+06 | 1.25 | 31.6 | 3.24 |

† Phenotypes are listed in order of decreasing volume in Design Space and cartoons of their topologies are shown ordered from right to left and top to bottom in **Figure 6(B and C)**. The elements of the system architecture are the following: C4 = cycle of four activations; C3 = cycle of three activations; L3 = link of three serial activations; L2 = link of two serial activations; L1 = link of one terminal activation; F3 = fan out of three activations; F2 = fan out of two activations; R2 = reciprocal pair of activations; A1 = autoactivation.
\* Although here is only one possibility with four auto activations (Case 121251), in most cases there are several phenotypes that are identical because of the symmetries in this system. Only a representative phenotype number is given; all other phenotypes with the same topology have the same global tolerances.
§ Global tolerance is the fold difference between the largest and smallest value of a parameter for which the qualitatively-distinct phenotype remains unchanged. There is a total of 28 parameters: 20 dissociation constants ( $K_{ij}$ values), four capacities for regulated expression ( $\rho_i = \alpha_{i\max} / \alpha_{i\min}$ values), and four maximum levels of expression ( $\gamma_i$ values). For any given phenotype, 12 of the dissociation constants are large enough to preclude binding, and beyond a minimum value of about 10 the capacities have no effect on the qualitative behavior. Thus, each phenotype is characterized by the quantitative values of the remaining 13 parameters. The minimum and maximum Global Tolerances are identified from the list for all parameters. A minimum estimate for the “volume” of a phenotype in System Design Space is given by the product of the Global Tolerances for the 13 parameters that characterize the phenotype. The (geometric) mean of the Global Tolerances is then the $p^{th}$ root of the volume, where $p = 13$ is the number of parameters that characterize the phenotype. The cooperativity $n = 2$ is assumed for all calculations.

Biological networks are generally recalcitrant to perturbation. Indeed, robustness of the decision-making circuity is one of the most important properties necessary to ensure proper differentiation in an inherently noisy intracellular and extracellular environment. The design space analysis provides a measure of global tolerances (Savageau et al, 2009) that define the size of disturbances (fold changes in the parameters and input variables of the system) that would cause a qualitative change of phenotype (i.e., loss of bistability). We further analyzed and ranked the global tolerance of the 19 different phenotypes for robustness in the face of large disturbances (**Table 1**). Those with the largest values have the largest region (“Volume”) in the 13-dimensional parameter space for which the bistable phenotype is preserved.

We next explored the consequences of a mutational event for these circuits - specifically, a loss or gain of a binding site for a given transcription factor, as could be produced by mutations that change the value of its dissociation constant. The change in circuitry exhibits a corresponding change in phenotype whose volume in parameter space is located adjacent to that of the original phenotype as “nearest” neighbors. For example, in the case of phenotype 336251 which is in the most robust class (**Figure 6B, Table 1)**, the nearest neighbors that still exhibit bistability are twelve phenotypes, 3 of which are shown in Figure 6C: 101876 (A1L3), 180001 (R2L2), 258126 (C3L1),. The aggregate “volume” in parameter space for these twelve bistable phenotypes is only 0.2% larger than that of the most robust phenotype alone. All of the other phenotypes are two or more mutational events removed from the most robust phenotype. Thus, from this analysis, the most likely circuit is predicted to be the four-gene cycle (**Figure 6B**) that is associated with phenotype 336251. A simulation of the bistable response to increasing VND7 induction for one of the genes in the cycle is shown in **Figure 7**.

**Figure 7.**
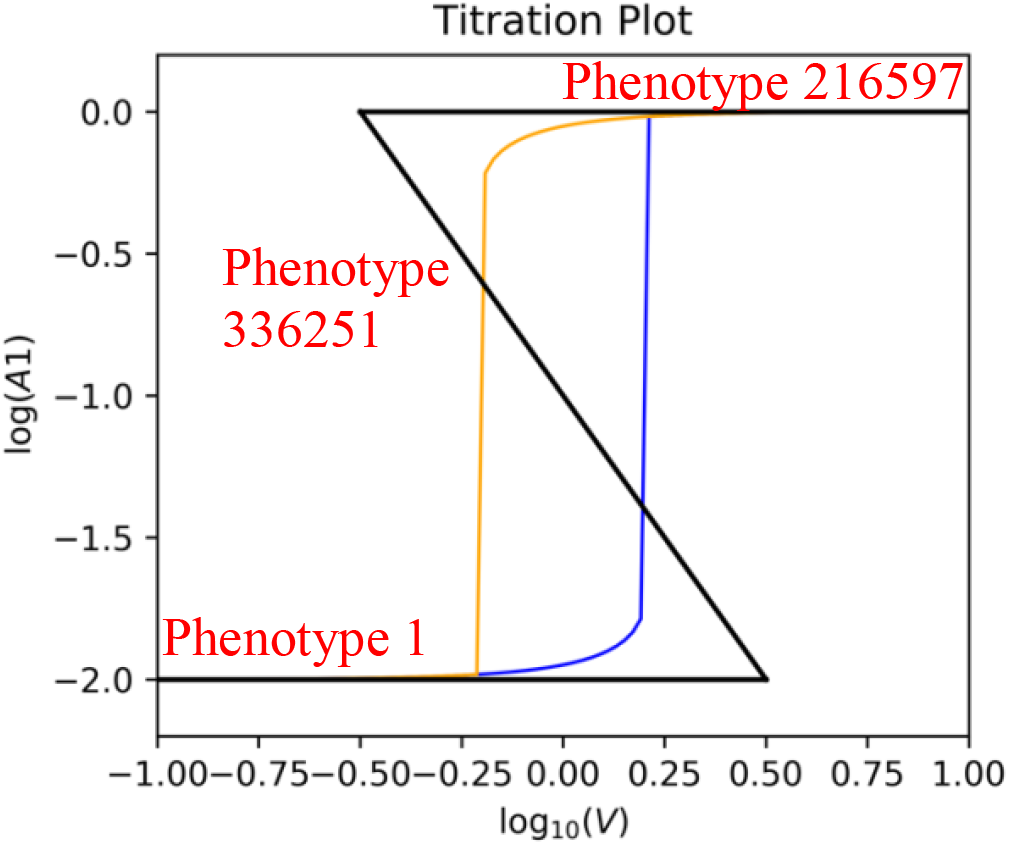
Bistable Induction by VND7 for the Four Class A Genes Linked in a Cycle. Only the response for the gene is shown; all three genes exhibit the same response because of the 4-fold rotational symmetry (**Figure 6B**). The three solid black lines are the predicted results from the System Design Space analysis: the lower stable states (phenotype number 1), the higher stable states (phenotype number 216597) and, in the case of the negatively inclined portion, the exponentially unstable states (phenotype number 336251). The orange and blue lines are the verified results obtained by simulating of the full system with the predicted values for the parameters [Eqns. 5-8]: (Blue) quasi-steady state titration with increasing values of V (VND7) and (Orange) quasi-steady state titration with decreasing values of V (VND7). Parameter values for both types of results were those predicted for the realization of phenotype number 336251 (**Table 1**), which exhibits the positive feedback necessary for exponential instability and the generation of a hysteretic bistable response: *n* =2; *K*_11_ = 100, *K*_12_= 0.316, *K*_13_= 100, *K*_14_= 100, *K*_1V_= 1.0; *K*_21_ = 100, *K*_22_= 100, *K*_23_= 0.316, *K*_24_= 100, *K*_2V_= 1.0; *K*_31_ = 100, *K*_32_= 100, *K*_33_= 100, *K*_34_= 0.316, *K*_3V_ = 1.0; *K*_41_= 0.316, *K*_42_= 100, *K*_43_ = 100, 1 *K*_44_ = 100, *K*_4V_ = 1.0; *γ*_1_= 1.0, *γ* _2_= 1.0, *γ* _3_= 1.0, *γ* _4_= 1.0; *ρ* _1_= 1.0, *ρ* _2_= 1.0, *ρ* _3_= 1.0, *ρ*_4_= 1.0. Note that relatively large values for dissociation constants are indicative of negligible binding.

## Discussion

Bistable hysteretic switches are a common mechanism by which cells commit to their final differentiated cell fate in animals. Bistable hysteresis requires both positive feedback and cooperativity. Our single-cell resolution data demonstrates cooperativity, although with the noisiness expected within a single cell population (**Figure 3C** and **Figure S3)**. Thus, these data support VND7 driving xylem cell differentiation via a bistable switch. Our whole root data further support VND7 acting as a bistable switch with hysteresis, although at the level of a whole population.

Sixty-five xylem cells out of 195 were observed upon induction within the whole root population (**Figure 3B**), which is less than the 100% observed in the elongation zone using confocal microscopy. The discrepancy is due to several factors including differences in the root spatiotemporal zone characterized as well as the developmental nature of xylem cells. Hair cells (trichoblasts), columella cells and “undefined” cells were not imaged in **Figures 2B** and **2C**. However, when these cell populations were removed from the single cell sequencing experiment demonstrated in **Figure 3B**, vascular cell types comprise 77.8% of induced cells in comparison to 14.6% in the controls. This is well within the 70-90% (log_2_ 6-6.5) range seen in Region 3 of **Figure 2**. Additionally, because xylem cells undergo programmed cell death we are unable to sequence the transcriptome of fully differentiated cells. Thus, we anticipate a decrease in the number of xylem cells identified in **Figure 3B** relative to **Figure 2C**.

The stochastic simulations using parameters provided from **Figure 2C** (**Figure S3)** demonstrated an interesting observation – with the noise observed at a single cell level, there is no way for a population of independent cells to show the results seen for the whole root. Using these simulations and the bistable case, if you average the trajectories for hundreds of cells, some fully ON (nearly all differentiated) and others fully OFF (a very small number of xylem cells), an average somewhere between ON and OFF would occur at the level of the whole population. This suggests that within the whole root induction experiment, there must be some mechanism that ‘coordinates’ the results for all the cells in the population to behave identically in the root elongation zone – fully un-induced or induced (**Figure 2C**). This is reminiscent of the theory of canalization (Sachs, 1968). Given that such behavior was not observed in the single cell profiling (all ON, or all OFF), we predict that disruption of the cell wall via protoplasting provides a hint that such a mechanism will likely involve cell-cell signaling.

Many of the TFs involved in xylem differentiation form a series of FFLs including both MYB46 and MYB83 (McCarthy et al., 2009; Taylor-Teeples et al., 2015). However, only *MYB46* expression closely mirrored the dynamics of *VND7* induction, and common VND7 and MYB46-direct targets were not enriched within the set of genes exhibiting bistable behavior. These data support our conclusion that the bistable switch observed is not caused by the VND7-MYB46-FFL. This reflects the observation that MYB46 is sufficient to regulate ectopic secondary cell wall deposition (including cellulose, xylan and lignin), but not xylem cell transdifferentiation (Zhong et al., 2007). Although MYB83 is also sufficient to only regulate ectopic secondary cell wall deposition like MYB46, it is only upon loss of both MYB83 and MYB46 function that a secondary cell wall phenotype is observed (McCarthy et al., 2009).

In addition to those genes identified to act in a bistable switch, a number of genes in four different classes (Classes D-G) were also identified showing a monostable-graded response upon *VND7* induction (**Figure 5A**). Here, expression of a set of hypothetical genes “Y” must respond to VND7 induction and provide the intermediary link to 37 monostable genes. Although our data demonstrates that VND7 regulates xylem transdifferentiation by a bistable switch with features of hysteresis, the enrichment of these “monostable, graded” genes suggest that VND7 may indeed have independent functions not associated with xylem cell transdifferentiation.

The regulatory logic which determines the bistable behavior of the 15 genes remains to be determined. The logic for the Class C gene (MYB46 direct target) is fairly straightforward in that it responds indirectly to VND7 induction with bistability likely either by autoactivating its own promoter or by receiving input from another bistable gene. The total number of possible phenotypes by which bistability could be achieved was beyond the scope of this study and deserves more experimentation, possibly by overexpression to determine if they are sufficient to drive xylem cell differentiation, or by CRIPSR-mediated editing of their promoters to abolish their regulation. In the case of autoregulation (the Class C gene) or the most robust four-gene circuit (the Class A genes, VND7 direct targets) we have a much more precise prediction of the most likely regulatory circuit that determines the bistable expression behavior. Similar experiments as those just described would be used to validate our predictions. In addition, the possibility of false positives or of false negatives could arise as a result of the noisiness of the single cell sequencing data. Furthermore, the possibility of reciprocal repression of Class A, B or C genes with a repressor could also give rise to such positive feedback. Single cell sequencing with increased depth coupled with additional experimentation should provide additional answers.

This VND7-mediated switch likely ensures that differentiation of xylem occurs rapidly as an “all or nothing” commitment. Given the importance of xylem cell fate commitment – in that, committing to differentiation results in cell death, this bistable relationship between *VND7* and xylem transdifferentiation buffers against fluctuations in *VND7* expression levels and requires *VND7* to first cross a threshold of expression before investing resources in secondary cell wall biosynthesis. Mapping developmental trajectories of other plant cell types will determine if this bistable hysteresis is an emergent property prevalent within only stem cells, like the cortex-endodermis initial cells, or also extends to cell types that undergo terminal differentiation, like xylem cells (Cruz-Ramirez et al., 2012). Further, it will be interesting to determine if the mechanisms for commitment to xylem cell development are similar to or different from those of animal cells, for which commitment to a terminally differentiated cell fate is common and totipotency is rare.

Finally, classification of cell identity from single cell RNA-seq profiles in this study relied on methods previously employed to characterize regeneration of a root stem cell niche (Efroni et al., 2015; Efroni et al., 2016). The clusters obtained by combining together cells from *VND7, MYB83* and *MYB46* overexpression lines revealed molecular differences that could not be completely differentiated given the general vascular ICI classifications and suggest potential new xylem cell subtypes. Further characterization of the marker genes that define these clusters would provide insight into the subtle differences by which VND7 and MYB46 regulate xylem cell differentiation. Enrichment of programmed cell death marker genes, genes co-expressed with *CESA4* and a mild enrichment of genes associated with lignin biosynthesis within the *VND7*-primary cluster suggests that the cells within this cluster represent the end of the xylem cell transdifferentiation developmental trajectory which includes bistability, programmed cell death, but not necessarily lignification (**Figure 4D**). The *MYB46*-primary cluster and associated enrichments show that MYB46 has the capacity to regulate some aspects of xylem cell differentiation or secondary cell wall synthesis distinctly from VND7 - specifically genes associated with hormone signaling and pattern differentiation (**Figure 4D**). The “mixed” cluster suggests that MYB46, MYB83 and VND7 coordinately regulate much earlier stages of vascular development, as this cluster is enriched in multiple cell identities in addition to the patterning genes and the auxin-responsive genes (**Figure 4D**). However, the enrichment of lignin biosynthesis genes is surprising and indicates that lignin biosynthesis may occur in vascular cells earlier than previously understood.

## Acknowledgements

GT was funded by an NSF predoctoral fellowship and AAUW Dissertation Completion Fellowship. GT, JRM and SB were partially funded by a Howard Hughes Medical Institute Faculty Scholar Fellowship. JRM was partially supported by a UC-MEXUS and CONACYT fellowship. MAS was supported in part by a grant from the US Public Health Service (RO1-GM30054). Single cell RNA sequencing work was supported by the Laboratory Directed Research and Development Program of Lawrence Berkeley National Laboratory and performed under U.S. Department of Energy Contract No. DE-AC02-05CH11231.

## Author Contributions

GMT & HV constructed and confirmed T3 homozygous estradiol inducible plant lines. GMT, JRM, MAS and SMB designed experiments to test VND7 as a bistable switch. GMT and DH conducted experiments confirming VND7 as a bistable switch in Figure 1. MAS wrote kinetic equations to model the system responses and analyzed global robustness of bistable phenotypes in the system design space. GMT, DH, SS and CNS conducted Drop-Seq experiments. GMT, SS, CNS, CJ, DED and SMB designed Drop-Seq experiments and analysis. All authors contributed to writing of the manuscript.

## Declaration of Interests

The authors declare no competing interests

## Material and Methods

### CONTACT FOR REAGENT AND RESOURCE SHARING

Further information and requests for resources and reagents should be directed to.

### Construction of Transgenic Lines

The *VND7* β–estradiol-inducible line was acquired from the TRANSPLANTA collection (NASC code: 2101676 TPT_1.71930.1C) (Coego et al., 2014). *MYB83* and *MYB46 TF*s were cloned in the Gateway vector derived from the β–estradiol-inducible vector PER8 (Zuo et al., 2000). Arabidopsis plants (Col-0) were floral dipped in *Agrobacterium tumefaciens* strain GV3101 (Bechtold and Pelletier, 1998). Transgenic plants were selected for on hygromycin and T3 homozygous plants were used for experiments. Increased expression of transgenics was confirmed with estradiol induction followed by qPCR (**Table S8**).

### Growth Conditions

Seeds were sterilized in a 50% bleach 10% tween solution and then stored at 4°C for 3 days. Sterilized seeds were germinated on nylon mesh (100 uM) filters on square MS petri dish plates and vertically grown at 22° C, 74% humidity in a 12 hr light cycle chamber at a 100-120 μmol m^−2^ s^−1^ light intensity. Seeds were plated in rows with approximately 100 plants per row and two rows per plate. Plates where randomized using a random block design where each tray contained an equal number of control and treatment plates. After 7 days of growth, plants were transferred by mesh filters to either MS plates buffered to pH 5.8 with KOH or MS containing estradiol (0.01, 0.05, 0.075, 0.1, 0.13, 0.15, 0.175, 0.2, 1, 2, 10, 20uM) plates and grown for an additional 24 hours (induction). All induction experiments were carried out at 7 am and collected at 7 am. Whole root samples were imaged, frozen for RNA extraction and qRT-PCR or protoplasted for Drop-seq.

## METHOD DETAILS

### Histology and Confocal Microscopy

Seven-day-old *VND7* over-expression roots were stained following a modified version of the ClearSee protocol (Kurihara et al., 2015; Ursache et al., 2018). Following 1 hour fixation with 4% paraformaldehyde in 1 x PBS, roots were transferred to the ClearSee solution overnight. Basic Fuchsin dye was used to mark lignified cells for identification of fully differentiated xylem. Roots were stained overnight in Basic Fuchsin followed by one ClearSee wash and an additional hour in the ClearSee solution. Roots were stained for an additional for 45 mins with Calcofluor White to stain the cellulose of the primary cell wall as well as the emergence of cellulose in secondary cell walls, allowing us to count the total number of cells in each image. Roots were visualized on the Zeiss LSM780 microscope: Basic Fuchsin: 550-561 nm excitation and 570-650 nm detection and Calcofluor White: 405 nm excitation and 425-475 nm detection. Z-stack images were collected from the start of the elongation zone.

### Confocal Laser Scanning Microscopy Analysis

The total number of cells was determined by the presence of an outline of a Calcofluor White stained cell. The Basic Fuchsin dye was used to mark lignified cells for identification of fully differentiated xylem. The percentage of cells experiencing trans-differentiation into protoxylem cells was determined by drawing a box around each cell in ImageJ and color coding based on presence or absence of a helical pattern (marked by Calcofluor or Fuchsin) as observed in **Figure 2B**. For each root, the area for each xylem marked cell was summed and divided by the total area of all cells. The percentage of cells experiencing trans-differentiation into protoxylem was determined for each estradiol concentration with 10-20 roots per concentration. Xylem differentiation was quantified for z-stacks throughout the entire root (**Table S1**).

### RNA extraction and qRT-PCR

Whole roots were separated from shoots and collected in bulk from each plate for RNA extraction. RNA for qRT-PCR was extracted using Trizol reagent (Life Technologies, USA) according to the manufacturer’s instructions. RNA was DNase treated prior to cDNA synthesis using M-MLV reverse transcriptase (Invitrogen). cDNA was used as a template for real-time quantitative PCR analysis. Each plate was considered an independent biological replicate with at least three biological replicates (same estradiol concentration & grown same day) per dilution series. Log_2_ Fold Change (FC) of VND7 varied 0.041 - 0.57 standard deviations (SD) among biological replicates. The largest variance was observed under lower induction of VND7 (2-4 FC) with the SD ranging from 0.08-0.57 SD, in mid-expression lines (5-7 FC) 0.041-0.064 SD. The 2^ΔΔCT^ method was used to measure relative transcript abundance (Schmittgen and Livak, 2008). Three technical replicates were carried out and averaged for each plate. VND7 expression was normalized to Actin to determine the delta CT and the delta delta CT was determined from uninduced roots grown on the same rack. Primers used for RT-PCR are listed in **Table S8**.

### Protoplasting and Drop-seq Assays

For Drop-seq runs, plants were grown as described above; the entire root was than separated and harvested for protoplasting (Benfey and Scheres, 2000; Birnbaum et al., 2003). A single Drop-seq sample consist of ten plates with ~200 roots per plate protoplasted in bulk as previously described. Individual cells were resuspended in Solution A containing: 600 mM Mannitol, 2 mM MgCl_2_*6H20, 0.10% BSA, 2 mM CaCl_2_*2H20, 2 mM MES Hydrate, 10 mM KCl and pH to 5.5 with 1 M Tris. Each sample of cells was filtered through a 40 μM Cell strainer (BD Falcon) into a 50 mL Falcon tube. Cells were then counted and diluted to 100-180 cells per μl immediately before Drop-seq (**Table S2**). All Drop-seq steps followed the standard protocol outlined by the McCarroll Lab (http://mccarrolllab.com/wp-content/uploads/2016/03/DropseqAlignmentCookbookv1.2Jan2016.pdf). The droplet diameter ranged from 0.78 - 1 nl, barcoded beads (Barcoded Beads SeqB; ChemGenes Corp., Wilmington, MA, USA) and cells were loaded at concentrations specified in **Table S2**, with a collection target of 1 mL according to standard Drop-seq protocol.

Drop-seq was run with a collection target of 1 mL according to standard Drop-seq protocol. cDNA was amplified using 13-17 cycles and double-purified in 0.6x volume of Agencourt AMPure XP beads. Tagmentation was carried out with 1200 pg of DNA as input. Samples were run on the Bioanalyzer (**Figure S8**) prior to NextSeq sequencing.

## QUANTIFICATION AND STATISTICAL ANALYSIS

### Alignment and Normalization of Drop-seq Data

All Drop-seq data was pre-processed and aligned and aligned to the TAIR10 *Arabidopsis* genome with STAR (Dobin et al., 2013) via the Drop-seq Tools v1.12 software (http://mccarrolllab.com/wp-content/uploads/2016/03/DropseqAlignmentCookbookv1.2Jan2016.pdf). Gene expression matrices were further processed and normalized using the Seurat R package, with a cutoff of at least 500 genes/cell and for genes seen in at least 3 cells (Satija et al., 2015). For each experiment the number of cells before and after processing are reported in **Table S2**. When scaling the data in Seurat for downstream analysis, cells with a high percentage of mitochondrial, chloroplast and protoplasting-induced genes were regressed out prior to clustering of cells (Birnbaum et al., 2003). Experiments in **Figure 4A** were merged using the MergeSeurat function, providing us more power to cluster cells but also the potential risk of clustering by experiment. Overall, we observed no large batch effects (**Figure 4A**).

### Determining Cell Identity

#### Clustering Analysis

Reducing data dimensionality and clustering cell types was performed as described by Andrews and Hemberg, 2018. Code is provided in github. To identify cell types via clustering we first reduced the data to the top 1,000 highly variable genes across cell types. We then further reduced the data dimensionality through principal component analysis (PCA). PCs 1-12 were used for further clustering as determined by the elbow of the PC standard deviations plot. Clusters were then determined from these PCs using the shared nearest neighbor modularity optimization based algorithm, Louvain clustering in FindClusters. To determine each cluster’s cell identity, the average expression value of each cluster was determined using Seurat’s AverageExpression function, requiring at least 25% of cells within the cluster to express each gene. These average expression profiles were then compared to ~100 predefined markers for cell identity in the Arabidopsis root and an ICI score calculated (Birnbaum and Kussell, 2011; Efroni et al., 2015). Tissues with significant p-values (< 0.01) for cell identity were used to define the corresponding clusters and cells within clusters. VND7, MYB83 and MYB46 were not among the 100 predefined markers for protoxylem identity induction experiments therefore did not bias assignment of cluster identity. To preserve the original data structure tSNE was used for plotting with cells color coded by cluster identity.

The number of cells in each DropSeq experiment before and after induction that could be assigned an individual cell identity given a significant ICI score are quantified in **Figure S9** and **Table S3**

The accuracy of cluster assignment was determined with the R Classification And REgression Training package; caret (Kuhn et al., 2008). First, the single cell data was resampled using cross validation, with 10 resampling iterations resulting in 10 training and 10 testing splits. The data was then trained using the support vector machine with linear kernel algorithm (svmLinear), as used by Seurat, to predict cluster identity given the expression of a subset of genes. Specifically, we focused on those with the strongest contribution to the first twelve principal components. After ten iterations of resampling the model prediction, accuracy and kappa were reported. When applied to the three protoxylem clusters in **Figure S7A-B** we observed 90% accuracy with clustering of only protoxylem cells and 99% accuracy in calling the protoxylem cluster when using all the data in **Figure 4A.**

Cluster accuracy was also determined by applying two clustering algorithms - Smart Local Moving (SLM) and Louvain clustering. When applied to the three protoxylem clusters of **Figure S7A-B** the two clustering algorithms agreed 100% of the time indicating that these three clusters indeed have distinct transcriptome profiles which is unbiased by the clustering algorithm used. The cells within these three clusters show 92% similarity to those seen in **Figure 4A**. Cluster 2 shows 100% similarity to cluster 10 of **Figure 4A** (the VND7 primary cluster), and cluster 1 showing the largest deviation from clusters of **Figure 4** with 90% similarity to cluster 7 (Mixed).

### Enrichment of Vascular Categories

Marker genes for each cluster were identified using Seurat with a log fold change threshold of plus two. Marker genes were compared globally using GO enrichment analysis in python (https://github.com/tanghaibao/goatools) or with predefined gene lists. In both cases a Fisher’s exact test was carried out comparing the study (marker genes of a cluster) to the population (all genes expressed in the single cell experiment), and p-values were adjusted with Benjamini-Hochberg. Gene lists were obtained from previously published work: lignin biosynthesis genes (Raes et al., 2003), CESA co-expressed genes (Persson et al., 2005), PCD (Olvera-Carrillo et al., 2015) and early vascular markers (Cano-Delgado et al., 2010). Genes related to auxin, cell size and patterning were determined by GO annotation.

### Identification of bistable behavior – the separation score

To determine VND7 downstream targets exhibiting bistable or monostable behavior we took advantage of the variance in *VND7* expression observed within single cell types. *VND7* expression was “binned” into three groups: low (0-2), middle (2-4) and high (4-6) VND7 normalized expression values. For the whole root data in **Figure 2C**, or the single cell data in **Figure 3C**, we measured the spread of xylem cell differentiation or of ICI score determination, respectively, by group these y-axes data into terciles or thirds. We next calculated a “separation score” for each of these measures. The separation score is the result of the 2/3 y-axis tercile value - 1/3 y-axis tercile value divided by the summed y-axis values. Our null hypothesis is that the mean separation score should be 1/3 if all data is evenly distributed.

In order to identify genes that respond to VND7 expression as a monostable or bistable “switch”, we identified genes from the single cell data which had a positive Pearson correlation with *VND7* (r^2^ > 0.5). Next, we divided cells into “xylem” cell populations (as determined using significant ICI scores) and non-xylem cells. Using custom scripts in R, we interrogated the correlated genes with VND7 expression by two means: a) compared expression of these genes in protoxylem induced cells against all uninduced cells on a bin-by-bin basis using the base R function *t.test()*. Next, we selected genes that were significantly different (p <= 0.05) in the medium and high bins but not in the low bin (p > 0.05). For further classification, the absolute difference between the mean expression of the middle and high bins was taken and the first (25%) quartile was selected (n=29 genes) (i.e., genes with minimal difference between the middle and high bins in the protoxylem induced cells, but significantly different between induced and uninduced). b) comparison between the middle and high bins using the 203 set of VND7 correlated genes, only in protoxylem induced cells; genes with no significant difference (t-test, p > 0.05) were selected (n=43 genes); The overlap between the sets in a) and b) was taken for further analysis as the set of potential bistable genes (n=15 genes). Conversely, the overlap between genes in the rest of the quartiles from **a)** and genes with significant difference (p < 0.05) in **b)** were selected as potential monostable genes. Enrichment of the clusters 5, 7 and 10 as well as the targets of VND7, MYB46 or VND7/MYB46 was done using the *fisher.test()* function in R with the monostable or bistable sets as defined above. GO enrichment was done using the gene list analysis tool in TAIR powered by PANTHER (www.arabidopsis.org, Mi et al 2012). AME (McLeay, 2010) was used for motif enrichment of the bistable gene set.

### Equations for generic switches

The generic model in **Figure S1A**, representing either bistable hysteretic or monostable graded switches, was represented by Eqns. (1-3) and used to generate the results in **Figure S1 B, C**.

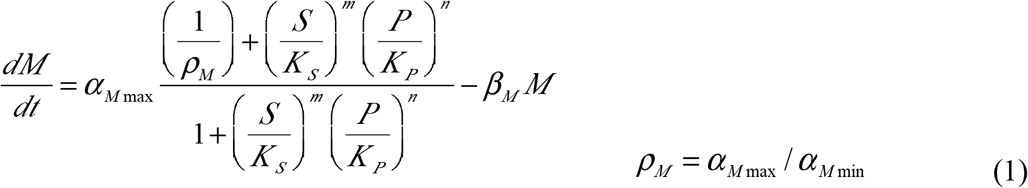

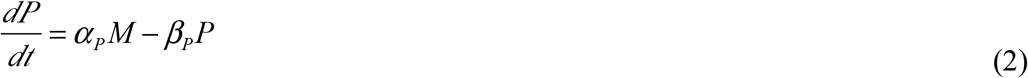

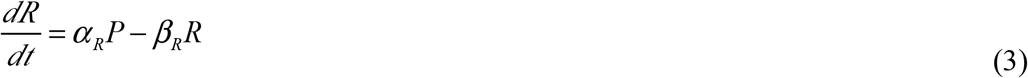

The messenger RNA (*M*) has a rate of synthesis that is modulated by the input signal (*S*), which is the transcription factor *TF*_1_, and the second transcription factor *TF*_2_ (*P*). The minimum and maximum rates of M synthesis are *α*_*Mmin*_ and *α*_*Mmax*_, the constants for half-maximum binding of *S* and *P* are *K*_*S*_ and *K*_*p*_, and the cooperativity (or effective Hill numbers) for binding *S* and *P* are the integers *m* and *n*. Messenger RNA (*M*) loss is governed by a first-order process with rate constant *β*_*M*_. The transcription factor *TF*_2_ (*P*) has a rate of synthesis proportional to the concentration of the message (*M*) with rate constant *α*_*p*_ and a rate of loss that is first-order with rate constant *β*_*p*_. The reporter (*R*) has a rate of synthesis proportional to the concentration of the transcription factor *TF*_2_ (*P*) with rate constant *α*_*R*_ and a rate of loss that is first-order with rate constant *β*_*R*_.

In steady state, these differential equations are solved for the transcription factor *P* as an implicit function of the stimulus *S* (Eqn. 4) and used to generate the results in **Figure 2C.**

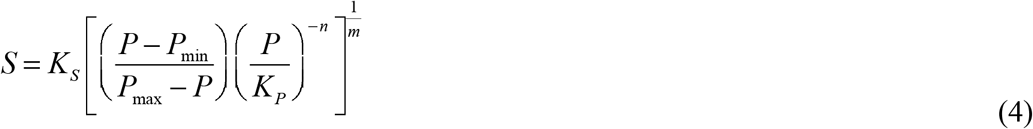

In steady state, all the derivatives are equal to zero, *M* in the second equation can be solved in terms of *P*, and this expression can be substituted into the first equation to obtain a single algebraic equation relating the steady state value of *P* to *S*. This algebraic equation has an implicit solution for the transcription factor *P* as a function of the stimulus *S* in which *P*_min_ = *α*_*M* min_*α*_*p*_/(*β*_*M*_*β*_*P*_) and *P*_max_=*α*_*M* max_ *α*_*p*_/(*β*_*M*_*β*_*p*_). The response (*R*) also is proportional to the transcription factor (*P*), from the third equation, and could be used in place of *P* in the resulting equation.

The parameters of this equation were used to fit the data in **Figure 2C** and used in the generation of **Figures S1D-F**.

### The Design Space Toolbox

The System Design Space Toolbox is a collection of novel analytical tools for the characterization and analysis of biochemical systems enabled by a rigorous mathematical definition of a system’s elemental phenotypes and conventional linear analysis of the transformed nonlinear systems (Savageau, et al., 2009; Lomnitz and Savageau, 2016). Among the tools used in this study are the following: (1) Exhaustive enumeration of a system’s phenotypic repertoire based on (a) system topology (interactions), (b) signs of the interactions (positive/negative), and (c) number of binding events in the interactions (small integer values). It is important to note that this phenotypic repertoire is obtained without the need to specify kinetic parameter values (binding constants, rate constants), which are typically unknown. (2) <u>Determination of input-output signal amplification factors, local parameter sensitivities and number of eigen values with positive real part</u> for each phenotype, again independent of kinetic parameter values. (3) <u>Prediction of kinetic parameter values for the realization of each phenotype</u> in the system’s repertoire. This involves the solution of a linear programming problem and requires no sampling or numerical estimation of parameter values. (4) <u>Deconstruction of parameter space into space filling regions</u> within which the values characterize a single qualitatively distinct phenotype. The boundaries are analytically determined, and intersecting regions define composite phenotypes (**Fig. 7**). (5) <u>Algebraic bifurcation analysis</u> (**Fig. 7**) differs from conventional bifurcation analysis, which is based on the numerical solution of differential equations and analytical continuation. For a comparison of these alternative approaches see (Lomnitz and Savageau, 2013; Valderrama-Gómez, et al., 2018). In addition to these and other novel tools, the toolbox includes traditional numerical simulation routines (**Fig. 7**). The following is a link to the downloadable software: http://bme.ucdavis.edu/savageaulab/software/.

### Determination of global tolerance

Global tolerance for a parameter or input variable of the system is the fold difference between the largest and smallest value of the parameter or input variable for which the qualitatively-distinct phenotype remains unchanged (Savageau et al, 2009). The minimum and maximum global tolerances for a given phenotype are identified from the list of global tolerances for all parameters and input variables that characterize the phenotype. A minimum estimate for the “volume” of a phenotype in System Design Space is given by the product of the global tolerances; namely, the 13 that characterize the phenotypes in Table 1. The (geometric) mean of the global tolerances is then the *p*^th^ root of the volume, where *p* = 13 is the number of parameters that characterize the phenotypes in Table 1.

This measure of global tolerance gives a minimal estimate; the actual “volume” in parameter space could be larger in cases with a very wide disparity between the longest and shortest aspect in non-orthogonal orientations (see Valderrama-Gómez, et al, 2018).

### Equations for regulatory models shown in Figure 6A

The generic model in **Figure 6A** was represented by Eqns. (5-8) and used to generate the results in **Table 1** and **Figure 7**.

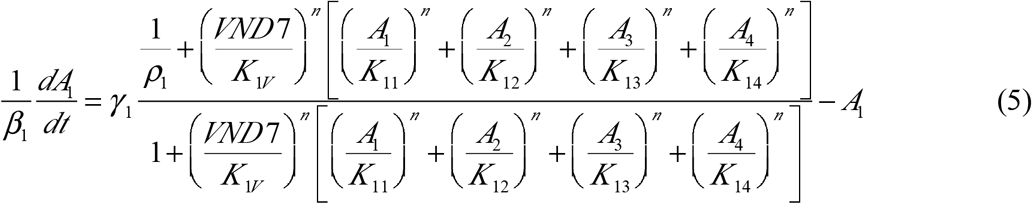

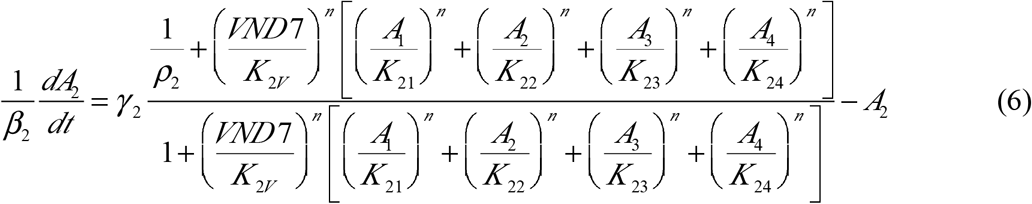

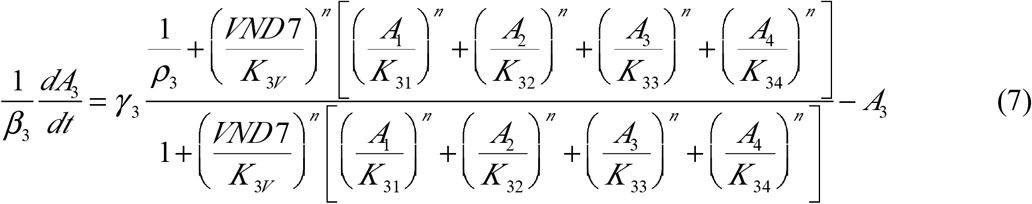

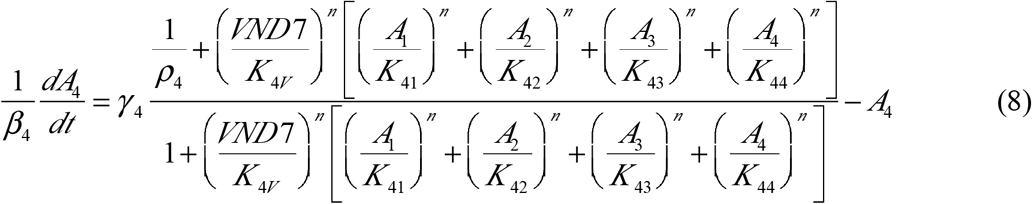

*VND* 7 is the concentration of the VND7 transcription factor; *A*_*i*_ is the concentration of the *i*^*th*^ transcription factor among the Class A genes. The *K* _*ij*_ parameters reflect the binding of the regulators to their promoter targets; *α*_*imax*_ the parameters are the maximum levels of transcription; the *β*_*i*_ parameters are first-order rate constants; and the capacity for regulation of a promoter is given by the ratio of maximum to minimum levels of expression *ρ*_*i*_ =*α*_*i*max_/ *α*_*i*min_. The concentration variables have been rescaled by dividing each equation by the corresponding *β*_*i*_ parameter; the maximum value for each concentration variable is then given by *γ*_*i*_ =*α* _*i max*_ /*β* _*i*_ and the minimum is given by γ _*i*_ / *ρ*_*i*_. The dynamics of mRNA are fast compared to that of protein; thus, protein concentrations are assumed to be proportional to their mRNA concentrations. The kinetic order *n* reflects the number of binding events (cooperativity) in the interactions (assumed to be the same in this case); there are no bistable phenotypes if *n < 2*, and the same bistable phenotypes are found if *n ≥ 2*. The cooperativity *n* = 2 is assumed for all calculations in **Table 1** and **Figure 7**.

## DATA AND SOFTWARE AVAILABILITY

Raw and processed Drop-seq data for VND7, MYB83 and MYB46 over-expression lines are deposited in GEO: <u>GSE114615. Code for determining cell identity from Drop-seq data and determining cell clusters were made</u> available on github (https://github.com/gturco/xylem_scRNA). <u>The Design Space Toolbox can be downloaded at http://bme.ucdavis.edu/savageaulab/software/</u>.

## KEY RESOURCES TABLE

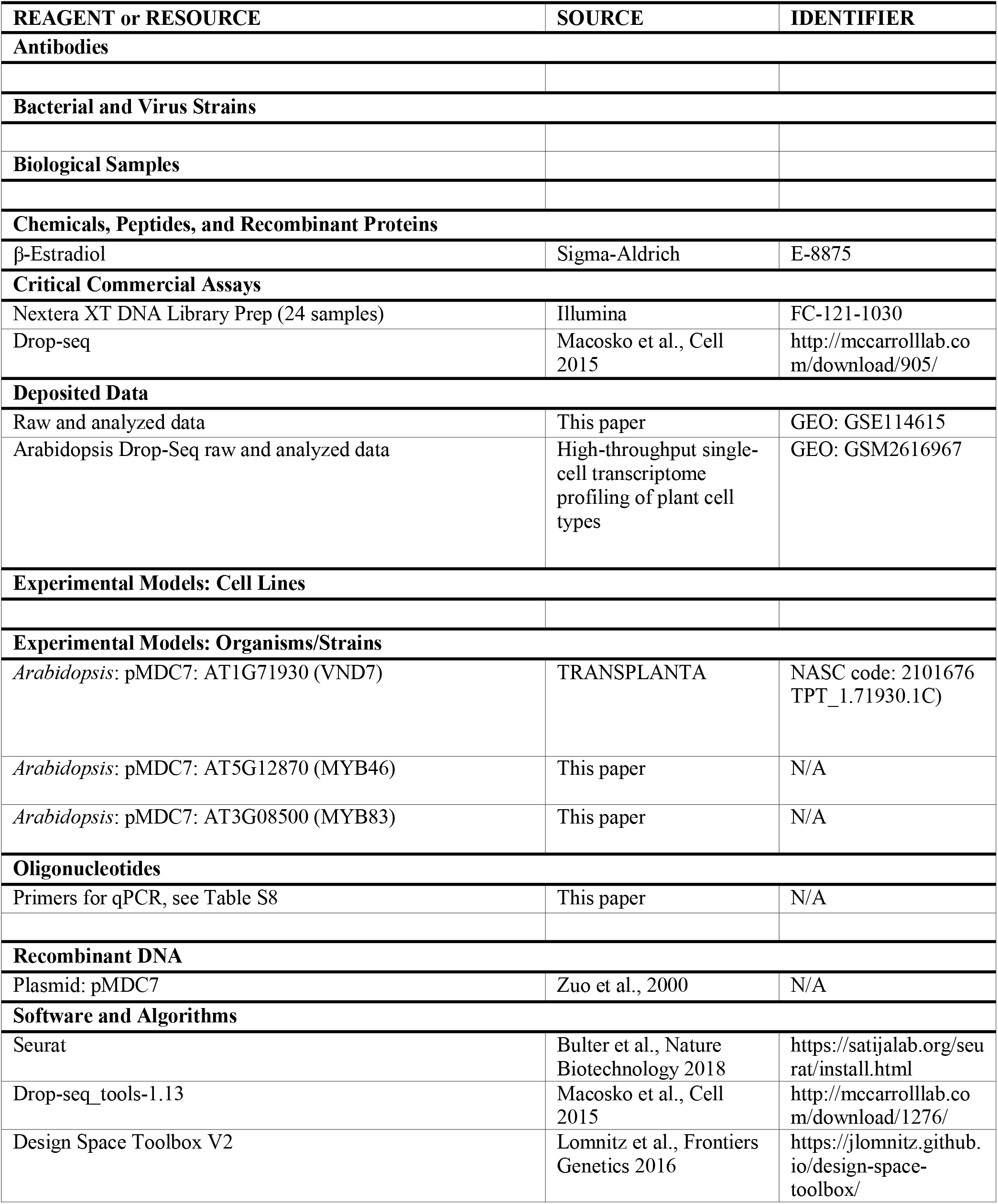
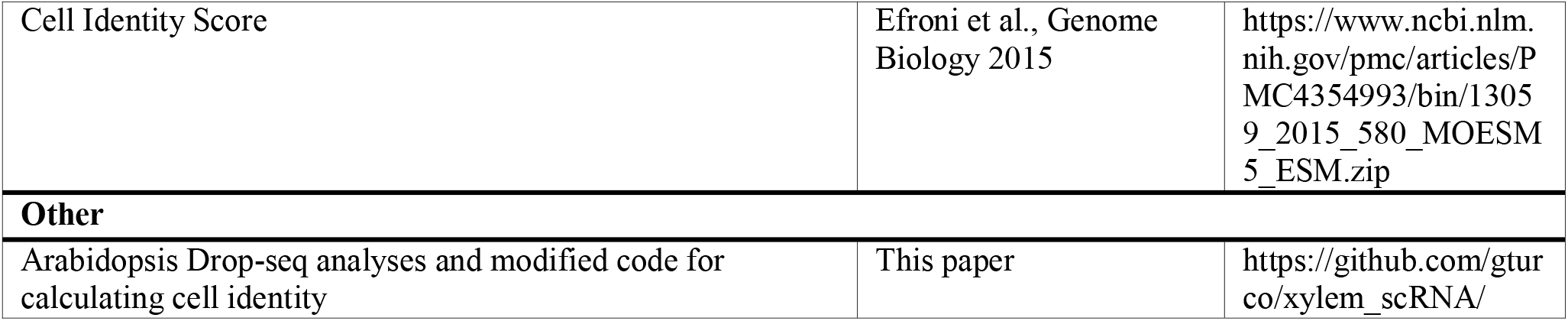

