## SupplementalFigures for "Molecular Mechanisms Driving Bistable Switch Behavior in Xylem Cell Differentiation"

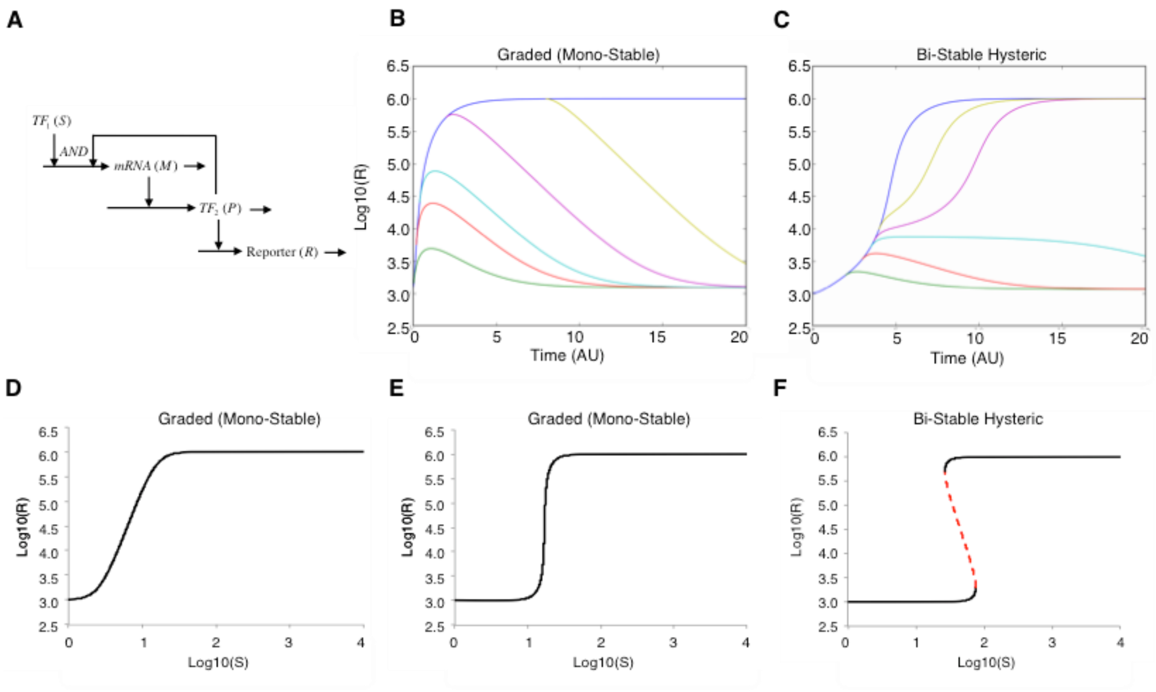


**Figure S1: Deterministic model capable of representing either bistable hysteretic or graded switching**. (**A**) System components: Transcription factor $TF_1$ (S), mRNA (M), transcription factor $TF_2$ (P), and reporter response (R). (**B,C**) Modeling the dynamics of the reporter (R) when naïve cells with the highest uninduced level of the developmental stimulus ($S = S_{Low}$) are treated with a fixed developmental stimulus sufficient for a full induction ($S = S_{High}$) for a fixed period of time ($\tau$). The response is followed as a function time following a return of the stimulus to the uninduced level of the naïve cells. (**B**) graded (monostable) or (**C**) bistable hysteretic responses, seen when both feedback and cooperativity are added to the model. (**D-F**) Steady state experimental strategy where naive cells are treated with different concentrations of the developmental stimulus (S) ranging from zero to a value sufficient for a full induction. Here the response is recorded as a function of the stimulus value rather than time. (**D**) A graded switch modeled when the network topology shown in **A** lacks positive feedback. (**E**) A graded response observed when positive feedback is added to the model but the feedback loop lacks cooperativity ($n \leq 1$). (**F**) Bistable switching observed in the model when both positive feedback and sufficient nonlinearity in the feedback loop are present. See methods for the mathematical equations. Common Parameters: (**B-F**) $V_{max}=90.6$, $V_{min}=6.1$, $K_S=10$, $K_P=260$, $\beta_M=1$, $\alpha_P=\beta_P=0.05$, m=4. Unique Parameters: (**B**) n=1, $S_{Low}=4$, $S_{High}=256$, $\tau=0.1,0.2,0.4,0.8,2.0,4.0,8.0$; (C) n=4, $S_{Low}=64$, $S_{High}=256$, $\tau=1.0,1.3,1.6,1.71,1.72,1.75,1.8$; (**D**) n=0; (**E**) n=1; (**F**) n=4.

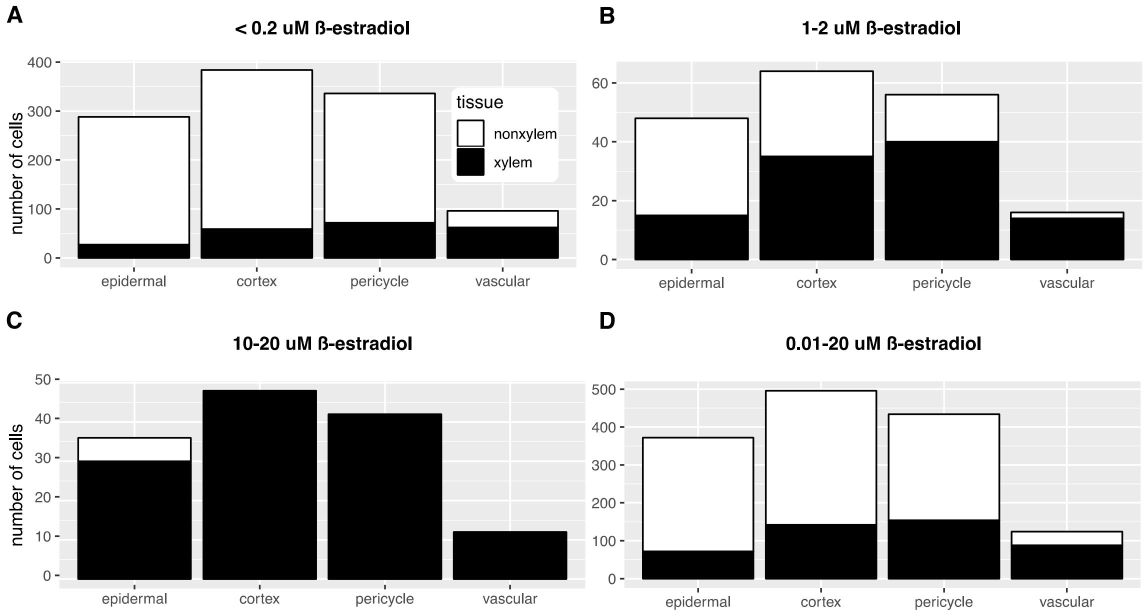


**Figure S2: Proportion of cells that have transdifferentated into xylem for each cell type given a range of VND7 induction.** Each cell type of the elongation zone, where cell types were inferred by positional information for multiple root images in *VND7* induced β-estradiol treated roots. In the "epidermal" column, the total number of cells in the epidermis position that are differentiated into xylem cell identity is indicated in black, while the number of cells that have the morphology of epidermal cells are indicated in white. Experiments are grouped into (A) . β-estradiol induction with <0.2 μM estradiol; (B) 1-2 μM estradiol; (C) 20 μM estradiol and (D) results summed over all estradiol concentrations.

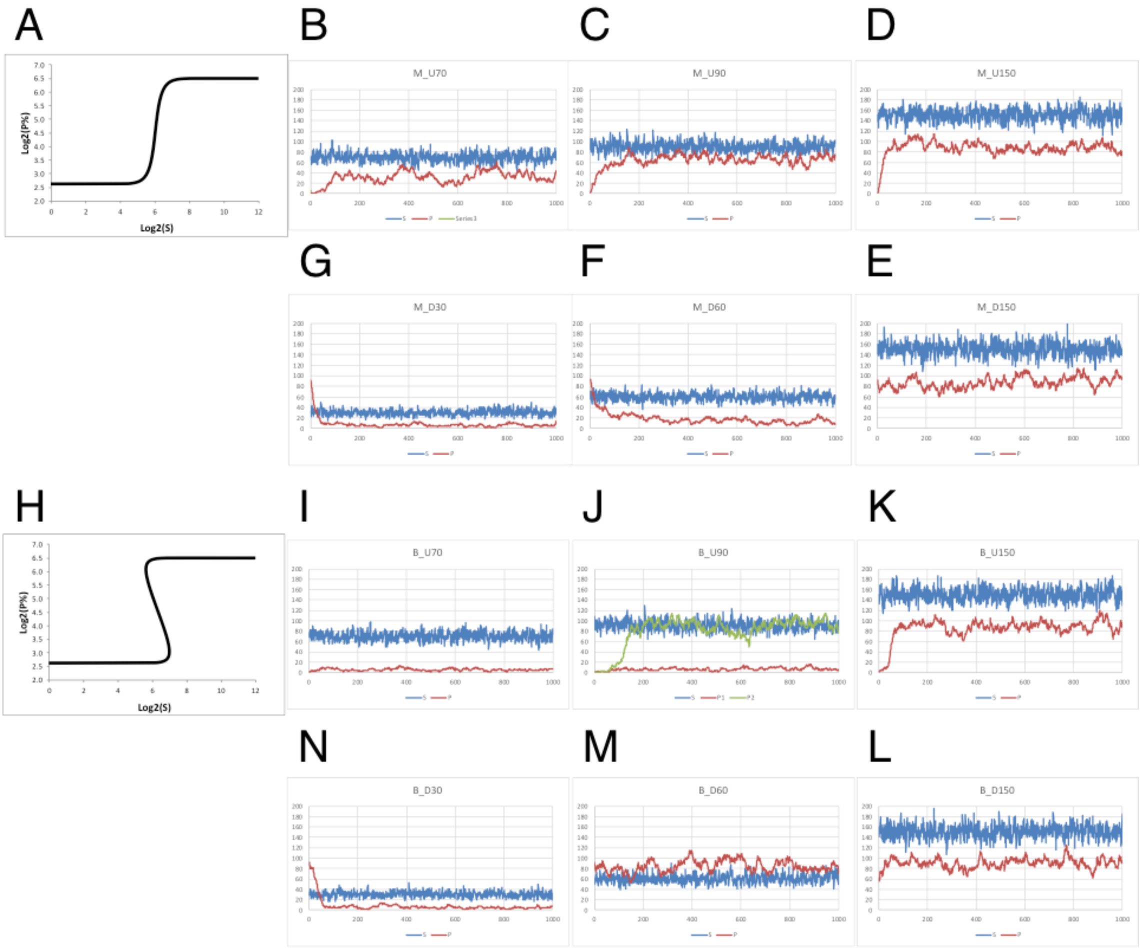


**Figure S3: Stochastic results for a mono-stable graded and bi-stable hysteretic System**. (**A-G**) Mono-stable system: The generic model is shown in Figure S1A**.** The deterministic results **(A)** as shown in Figure S1E for a mono-stable response. **(B-D)** Temporal behavior of the stimulus (blue trajectories) and response (red trajectories) for increasing levels of the mean stimulus. At time zero, the response is set equal to the minimum level of expression and the stimulus is *increased* from S=0 to **(B)** S=70, **(C)** S=90, and **(D)** S=150. **(E-G)** Temporal behavior of the stimulus (blue trajectories) and response (red trajectories) for decreasing levels of the mean stimulus. At time zero, the response is set equal to the maximum level of expression and the stimulus is *decreased* from S=200 to **(E)** S=150, **(F)** S=60, and **(G)** S=30. (**H-N**) Bi-stable system: The generic model is shown in Figure S1A**.** The deterministic results **(H)** as shown in Figure S1F for a bi-stable response. **(I-K)** Temporal behavior of the stimulus (blue trajectories) and response (red and green trajectories) for increasing levels of the mean stimulus. At time zero, the response is set equal to the minimum level of expression and the stimulus is *increased* from S=0 to **(I)** S=70, **(J)** S=90, and **(K)** S=150. **(L-N)** Temporal behavior of the stimulus (blue trajectories) and response (red trajectories) for decreasing levels of the mean stimulus. At time zero, the response is set equal to the maximum level of expression and the stimulus is *decreased* from S=200 to **(L)** S=150, **(M)** S=60, and **(N)** S=30. Only the bi-stable system exhibits the mean hysteresis observed in the experimental data of **Figure 2C**. See methods for the mathematical equations. Common

Parameters: $V_{max}$=90.6, $V_{min}$=6.1, $K_S$=10, $K_P$=260, $\beta_M$=1, $\alpha_P$=$\beta_P$=0.05, m=4. Unique Parameters: **(A-G)** n=1 and **(H-N)** n=4. Stochastic simulations were performed with the StochSS software (Drawert, et al., 2016).

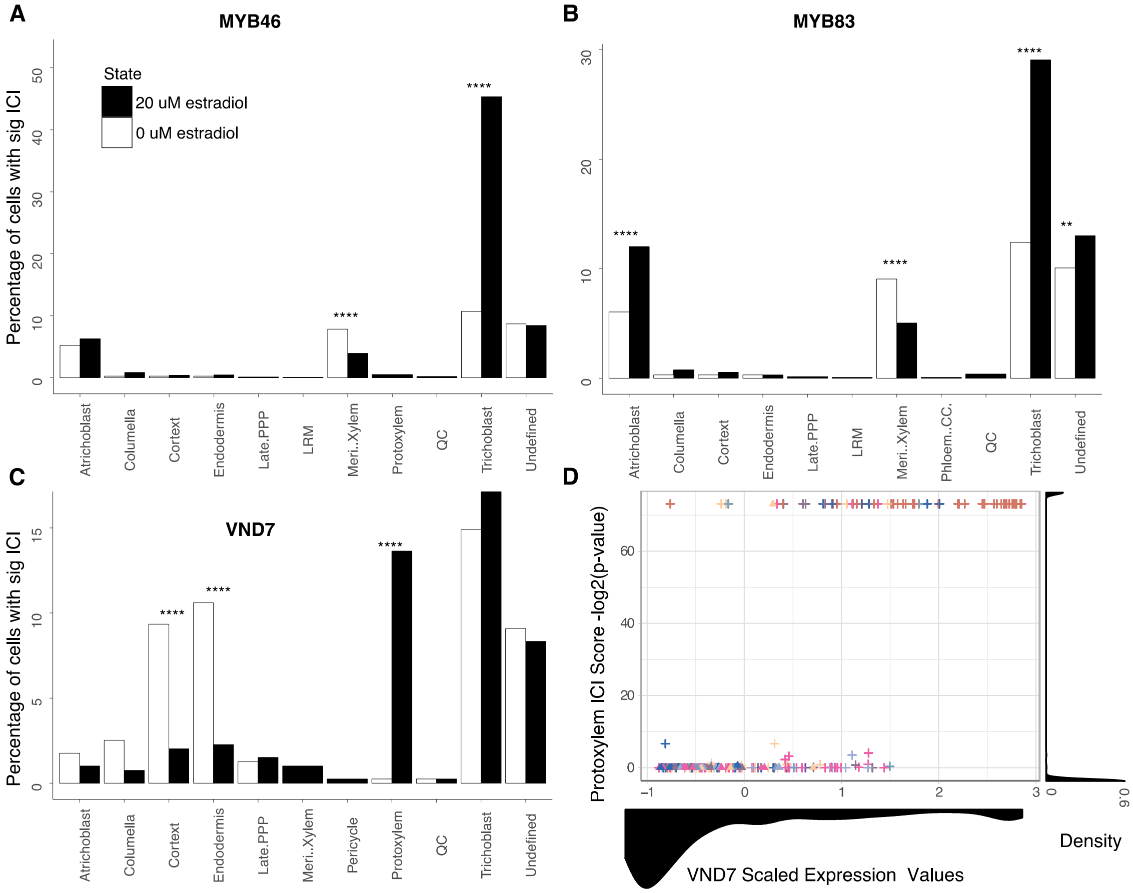


**Figure S4: Quantification of cell type/identity in MYB46 and MYB83 overexpression experiments.** Quantification of cell identity determined from ICI scores for individual cells in the induced population (20 μM β-estradiol), compared to the percentage of cells with that identity in the un-induced population, (0 μM β-estradiol). **** represents an adjusted p-value $\leq$ 0.0001, *** p-value $\leq$ 0.001, ** p-value $\leq$ 0.01, * p-value $\leq$ 0.05 as determined by a Fisher's-Exact test with Benjamini-Hochberg correction for **(A)** VND7, **(B)** MYB46 and **(C)** MYB83 over-expression (β-estradiol induction) experiments. **(D)** Cell density is plotted for each axis of related **Figure 3C**. The graph along the x-axis represents the density of cells at varying VND7 expression levels while the y-axis plots density of cells with a particular protoxylem ICI score.

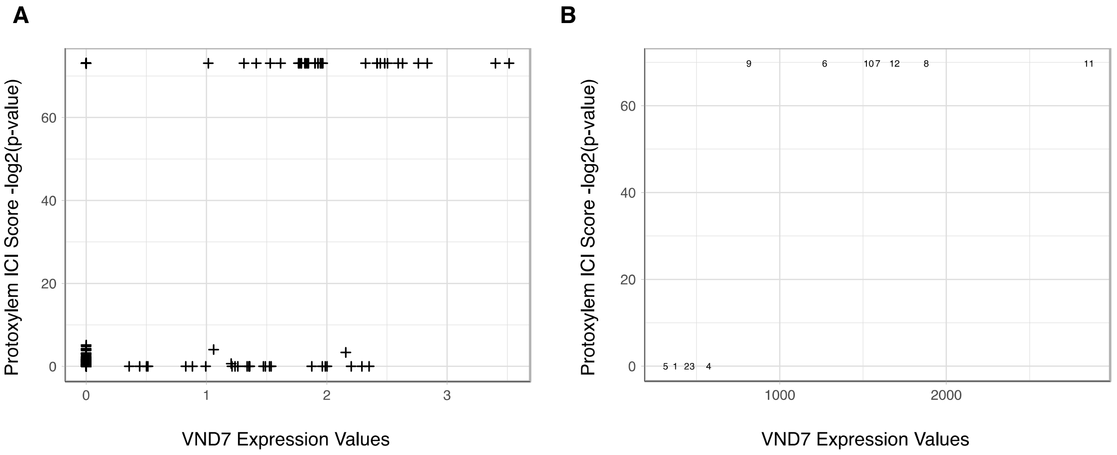


**Figure S5: Evidence for a bistable transcriptional switch in native cells. (A)** Log2 normalized VND7 expression values from ~12,000 wild type Arabidopsis root cells and their corresponding xylem ICI -$\log_2$(p-value). Each X represents an individual cell data taken from (Shulse et al, under revision, Nature Plants). **(B)** Arabidopsis root microarray imputed data for xylem developmental time. Spatiotemporal expression values are imputed according to methods described by (Cartwright et al., Bioinformatics, 2009). Here each point/number represents the root section number where lower numbers are closer to the root tip. VND7 imputed expression values are plotted on the x-axis for each xylem section and the corresponding xylem ICI -$\log_2$(p-value) on the y-axis.

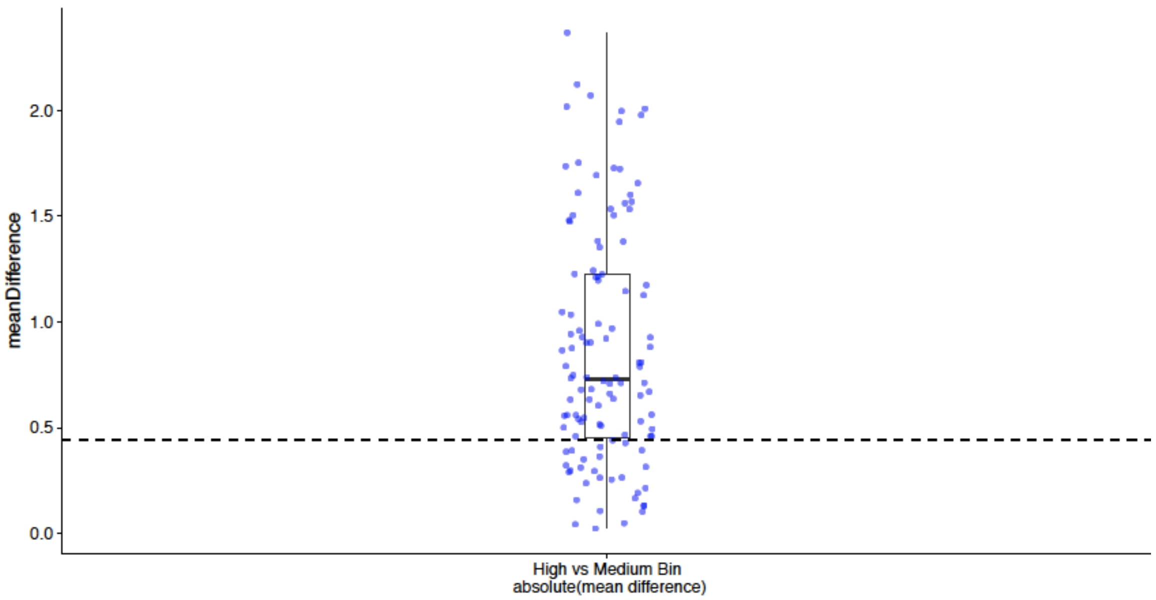


**Figure S6: Identification of bistable genes.** Boxplot of the mean difference between the high and medium bin for each gene that is correlated with VND7 at r=0.5. The dashed line indicates the bottom quartile of genes which are the candidates for bistability.

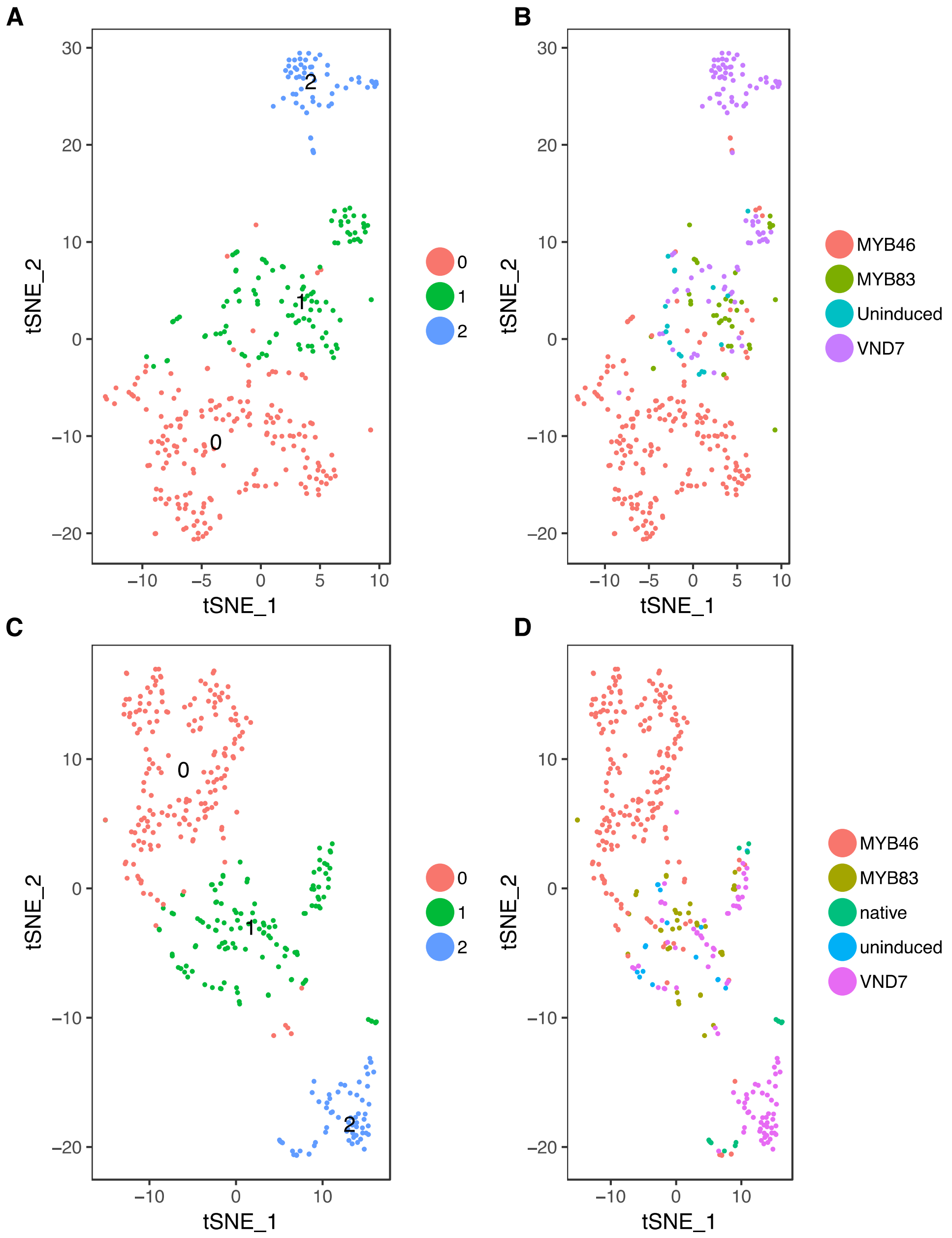


**Figure S7: Protoxylem cell clusters of Figure 4A.** tSNE plot of three protoxylem clusters depicted in **Figure 4A**. Here clusters where re-clustered with Louvain using only cells identified in **Figure 4A** as belonging to protoxylem or protoxylem-like clusters. **(A)** Each dot represents an individual cell colored by protoxylem cluster identity. **(B)** Each dot represents an individual cell colored by experimental origin of the cell. **(C)** Re-clustering of three protoxylem clusters with 18 wild-type (native) xylem cells from (Shulse et al., 2018), cells are colored by cluster identity and **(D)** experimental origin.

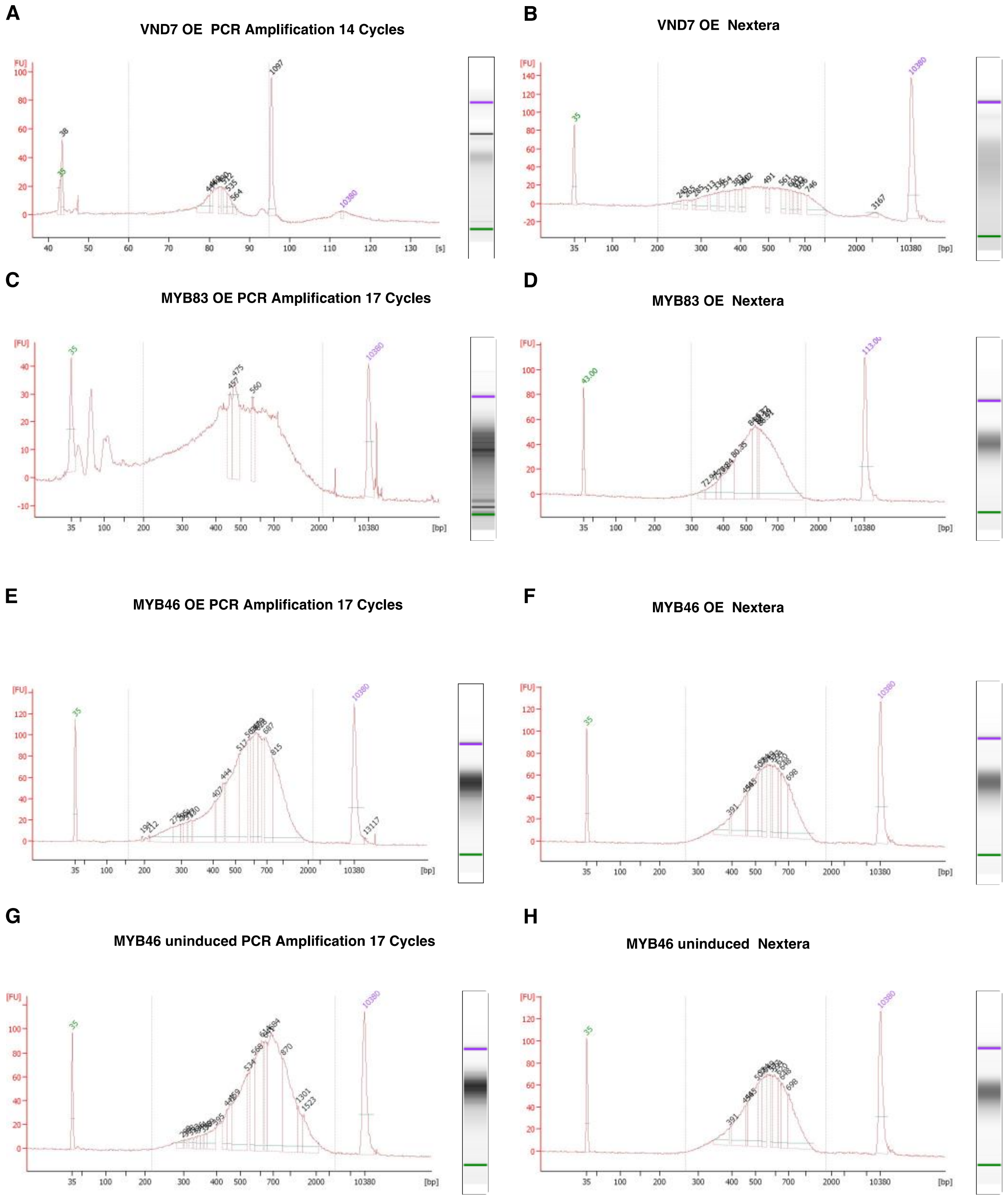


**Figure S8: Representative bioanalyzer results for Drop-Seq experiments. (A)** DNA from *VND7* induced roots after 14 cycles of PCR amplification **(B)** after Nextera **(C)** DNA from *MYB83* induced roots after 17 cycles of PCR amplification **(D)** and Nextera. **(E)** DNA from *MYB46* induced roots after 17 cycles of PCR amplification **(F)** and Nextera. **(G)** DNA from *MYB46* uninduced roots (representative of control roots) after 17 cycles of PCR amplification **(H)** and Nextera.

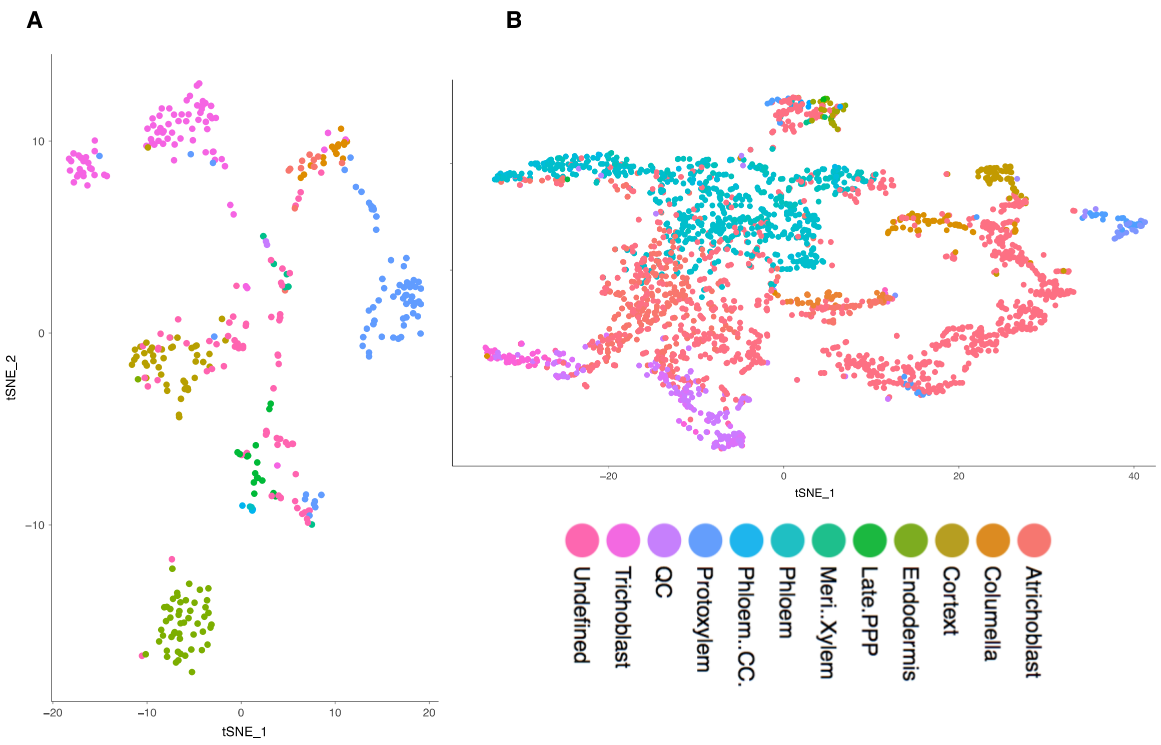


**Figure S9: Single cell clusters colored by individual identity.** tSNE plots from **(A) Figure 2A** and **(B) Figure 4A** where each dot represents an individual cell colored by its individually determined cell identity instead of previously shown cluster identity. In **B** there are more undefined cells because the number of cell numbers increased resulting in a lower sequencing depth per cell.

**Table S6**: Primers for qRT-PCR and semiquant RT-PCR

| **GeneID** | **Name** | **Forward Primer** | **Reverse Primer** | **Amplicon length** |
|---|---|---|---|---|
| AT3G08500 | MYB83 | AGCAACAACCTAAACTTACAGGCC | CCCAACACATTCCCTACCTT | 92 bp |
| AT5G12870 | MYB46 | GAATGTGAAGAAGGTGATTGGTACA | CGAAGGAACCTCAGTGTTCATCA | 130 bp |
| AT1G71930 | VND7 | CGAAGGAACCTCAGTGTTCATCA | GGGAAGCATCCAAGAGAA | 110 bp |
| AT4G18780 | CESA8 | TGATTGCTCGCCTTCTCTTT | CGGTTGATCATCTTGGCTCT | 207 bp |
| AT5G67210 | IRX15-L | ACGGGCGTACAGTTTTCATC | TTCTGCACTGGACGACACTC | 172 bp |

| | | | | |
|---|---|---|---|---|
| AT5G03260 | LAC11 | CACGGCTAACCTTGGAACAT | ATCAGCCCTGAATCTGATGG | 271 bp |
| AT4G05320 | UBQ10 | GGCCTTGTATAATCCCTGAT | AAAGAGATAACAGGAACGG | 61 bp |
| AT1G13320 | PP2AA3 | GGGCAATGCAGCATATAG | CCAAACACAATTCGTTGCTG | 389 bp |
| AT3G18780 | ACT2 | ACGGTAACATTGTGCTCAG | TTGGAGATCCACATCTGCT | 197 bp |
| AT5G44030 | CESA4 | AAATGGACAACCCTTCTCA | GAACACCCAAATCGGAAA | 399 bp |
| AT5G62380 | VND6 | CATCATCTCTCTGAAGAGCA | CGTGTGTGTATTTTGAGCC | 101 bp |
| Estradiol XVE protein | XVE | GTGCCTGGCTAGAGATCCTG | GATGTGGGAGAGGATGAGGA | 401 bp |
